# Graph-structured populations elucidate the role of deleterious mutations in long-term evolution

**DOI:** 10.1101/2024.07.23.604724

**Authors:** Nikhil Sharma, Suman G. Das, Joachim Krug, Arne Traulsen

**Affiliations:** Department of Theoretical Biology, Max Planck Institute for Evolutionary Biology, August-Thienemann-Str. 2, 24306 Plön, Germany; Institut für Ökologie und Evolution, Universität Bern, 3012 Bern, Switzerland; Swiss Institute of Bioinformatics, 1015 Lausanne, Switzerland; Institute for Biological Physics, University of Cologne, D-50937 Cologne, Germany

## Abstract

Birth-death models have long been employed to understand the interplay of genetic drift and natural selection. While well-mixed populations remain unaffected by the choice of replacement rules, the evolutionary outcomes in spatially structured populations are strongly impacted by this choice. Moving parent individuals to vacant sites gives rise to new update rules, leading to new fixation categories for spatial graphs. We discover a new category of graphs, amplifiers of fixation, where a structure has a higher probability of fixation for mutants than the well-mixed population, regardless of their fitness value. Under death-Birth updating with parents moving to vacant sites, the star graph is an amplifier of fixation. For very large population sizes, the probability to fix deleterious mutants on the star graph converges to a non-zero value, in contrast to the result from well-mixed populations where the probability goes to zero. Additionally, most random graphs are amplifiers of fixation for death-Birth updating, with parent individuals replacing dead individuals. Conversely, most random graphs are suppressors of fixation− graphs with lower fixation probability for mutants regardless of their fitnesses− for Birth-death updating with offspring replacing dead individuals. When subjected to long-term evolution, amplifiers of fixation, despite being more efficient at fixing beneficial mutants, attain lower fitness than the well-mixed population, whereas suppressors attain higher fitness despite their inferior ability to fix beneficial mutants. These surprising findings can be explained by their deleterious mutant regime. Therefore, the deleterious mutant regime can be as crucial as the beneficial mutant regime for adaptive evolution.

## I. INTRODUCTION

Spatial structure can substantially impact the evolution of a population [1–11]. Understanding the role of spatial structure in evolutionary biology is crucial and demands moving beyond the commonly assumed well-mixed populations [12]. Evolutionary graph theory provides a platform where structured populations are modelled as graphs [13], with each node representing an asexually reproducing individual and the links defining its interaction neighbourhood. In this framework, a complete graph represents a well-mixed population where each node interacts with every other node with equal propensity.

Fixation probability and fixation time are two key observables in evolutionary graph theory. The fixation probability of a mutant is the probability for the mutant to take over a population of wild-types [14–18], while the (conditional) fixation time represents the duration it takes for this process to complete [19–22]. The fixation time of a mutant is a random variable with a specific distribution, and the quantity of interest generally is the average fixation time [23].

The fixation probability is an important quantity in evolutionary biology, because it determines the rate of evolution [24–26]. During a fixation event, two evolutionary forces are at play – natural selection and genetic drift – and spatial structure can modulate the strength of these forces [27]. With the well-mixed population serving as the reference, graph structures that amplify the strength of selection are termed amplifiers of selection (*AoS*), while those that suppress it are referred to as suppressors of selection (*SoS*) [13, 15]. An *AoS* is a graph that has a higher probability of fixing beneficial mutants and a lower probability of fixing deleterious mutants compared to the complete graph. On the other hand, a *SoS* is a graph that has a lower probability of fixing beneficial mutants and a higher probability of fixing deleterious mutants.

In general, graphs can be weighted and directed [28–30]. This means that an individual may not interact with its neighbor as strongly as the neighbor interacts with the focal individual. In this work, we focus on unweighted and undirected graphs. The precise form of the interactions among individuals is determined by an update rule. The commonly studied update rules in evolutionary graph theory are the Moran Birth-death (Bd) and the Moran death-Birth (dB) updates [31–33]. The shorthand Bd implies that the birth event precedes the death event. The uppercase B indicates that selection operates during the birth event, while the lowercase d represents the neutral nature of the death event. This offers various choices for the birth-death updates [34]. The fixation probability of a mutant on a graph depends crucially on the update rule [35]. In [33] it was found that for Bd updating, most small random graphs are *AoS*, whereas under dB updating, most of the random graphs are *SoS*.

However, not only the update rule, but also the node where the mutant initially appears substantially affects the fixation probability. Mutant initialisation schemes determine the likelihood for a node to be initialised with the mutant. Two popular schemes are uniform mutant initialisation and temperature mutant initialisation [36]. Under uniform mutant initialisation, every node is equally likely to be initialised with the mutant. For temperature initialisation, the initial mutant is more likely to appear on nodes with higher turnover rates, that is, with a larger number of links. The star graph is an *AoS* under Moran Bd updating with uniform mutant initialisation. However, for temperature initialisation, the star graph is a suppressor of fixation (*SoF*) −a graph with lower probability of fixing a mutant than the well-mixed population, regardless of the mutant’s fitness.

Recently, some studies in evolutionary graph theory went beyond the fixation time scales and explored the state of mutation-selection balance that emerges at long times [37–39]. When the uniformly initialised star graph (an *AoS*) was subjected to long-term mutation-selection dynamics, it achieved a higher average steady-state fitness compared to the complete graph [40]. This outcome was anticipated because an *AoS* is more efficient at fixing beneficial mutants and preventing the fixation of deleterious mutants. Surprisingly, however, the temperature initialised star graph, despite being a *SoF*, not only attained a higher fitness than the complete graph, but also equally high fitness as the uniform initialised star graph. This result can be explained by the ability of the temperature initialised star graph to efficiently reject deleterious mutants, compensating for its inability to fix beneficial mutants. To our knowledge, this is the only known example so far where the deleterious mutant regime has the potential to influence long-term evolution, and it raises a number of questions. How common are *SoF* for temperature initialised Bd updating? Do all of *SoF* attain higher fitness than the complete graph, despite having lower probabilities of fixing advantageous mutants? What about dB updating? Does the deleterious mutant regime play any significant role for dB long-term dynamics as well? We address all of these questions here.

The structure of this paper is as follows. We begin by establishing a connection between update rules and mutant initialisation schemes. We show that these schemes naturally arise from the choice of individuals moving to vacant nodes – either the parent-type offspring or the mutant offspring – and therefore do not need to be separately specified. Subsequently we study the star graph and Erdős-Rényi random graphs at short-term fixation time scales, considering various update rules. Notably, we observe that the star graph acts as an amplifier of fixation (*AoF*) under temperature initialised dB updating with a higher probability of fixing mutants compared to the complete graph, regardless of the fitness value. Similarly, we find that most of the small random graphs are *AoF* under temperature initialised dB updating and *SoF* under temperature initialised Bd updating. Additionally, we study the star graph and random graphs under long-term mutation-selection dynamics. Surprisingly, despite being *SoF*, most of the random graphs achieve higher fitness than the complete graph for temperature initialised Bd updating, whereas most of the random graphs attain lower fitness for temperature initialised dB updating despite being *AoF*.

## II. UPDATE MECHANISMS IN GRAPH STRUCTURED POPULATIONS

When working with structured populations, the update rule substantially affects the dynamics. An evolutionary update rule determines not only the order of the birth and death events and the choice of event(s) where selection operates, but also the process by which new mutations appear in the population.

A mutant initialisation scheme *ℐ* denotes the probability distribution with which an initial mutant appears on a node, *p* = (*p*_0_, *p*_1_, · · · *p*_*N*−1_). For example, in the uniform mutant initialisation scheme *𝒰*, we have *p*_*i*_ = 1*/N* for all nodes *i*, i.e., a mutant appears in every node with the same probability. Similarly, for the temperature initialisation scheme *𝒯*, the probability for a node to receive an initial mutant is proportional to the temperature (sum of incoming/outgoing weight) of the node. We assume that mutation is coupled to reproduction.

As an example for temperature initialisation, let us focus on the Moran Birth-death (Bd) update rule with the offspring moving to another site [36]. First an individual is selected with probability proportional to its fitness to give birth to an offspring. Thus fitness is equivalent to the reproduction rate of an individual. After reproduction, the offspring either resembles its parent with probability 1− *µ* or is a mutant with probability *µ*. Then the offspring takes over the node of a random neighboring individual chosen to die. Therefore, in a population where every individual has the same fitness, the first mutant is more likely to appear on nodes with higher in-degree. To be specific, the probability *p*_*i*_ that a mutant appears in a node *i* is proportional to its in-temperature 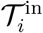,

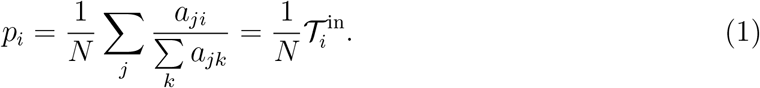

Here, *a*_*lm*_ is an element of the adjacency matrix **A** with value equal to 1 if there is link directed from node *l* to *m* and 0 otherwise. **A** is a symmetric matrix for undirected graphs. For further examples for this update mechanism, see also Refs. [41, 42].

So far, we assumed that the offspring moves to a neighboring node while the parent remains at its position, see Fig. 1 A. To denote this, we use the shorthand Bd^*o*^ where the superscript indicates that the offspring moves to a vacant site. With the same assumption, for death-Birth updating dB^*o*^, the mutant initialisation is uniform: The probability that an initial mutant appears in node *i* under dB^*o*^ updating is equal to 1*/N*.

**FIG. 1.**
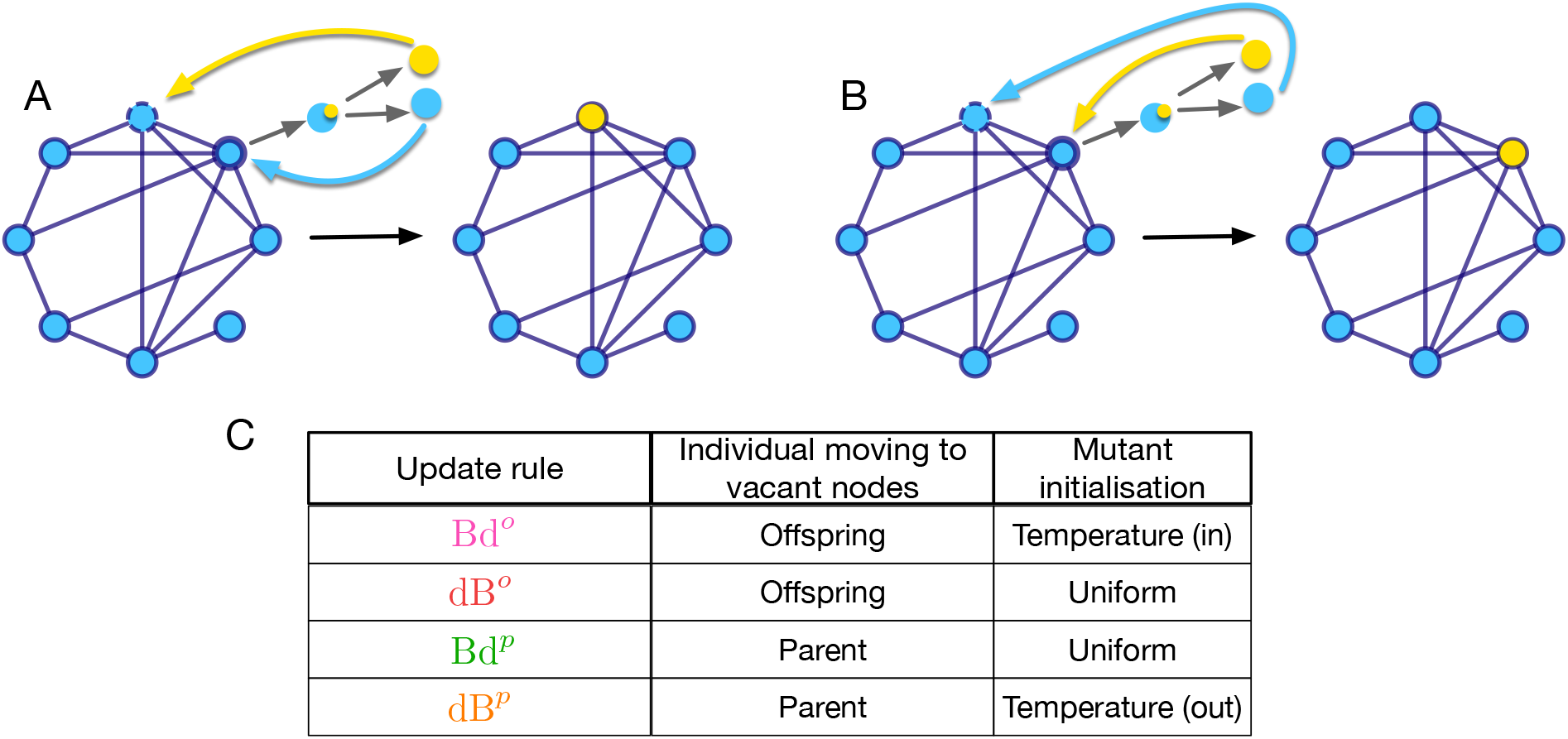
Type of individual moving to vacant sites and mutant initialisation schemes. The mutant initialisation scheme, the likelihood that the initial mutant appears on a given node in a homogenous fitness background, is fully determined by the evolutionary update rule, provided mutations are coupled to reproduction and the choice of individual that moves to the vacant node is specified. This is shown in panels A and B, where wild-type individuals are shown in blue and the mutant in yellow. The individual chosen for birth is marked by a thick solid circle while the one chosen for death is marked by a dashed circle. In case of mutation, one of the daughter cells is a mutant (yellow type) and the other one resembles the mother (blue type). In panel A, the mutant offspring moves to the vacant site. Throughout the paper, this is referred to as the offspring moving rule. For Bd updating, the corresponding initialisation scheme Bd^*o*^ is temperature initialised, and for dB updating, dB^*o*^, it is uniformly initialised. Similarly, in panel B the parent-type offspring moves to the vacant node while the mutant offspring stays at the birth site. We call this the parent moving rule. The corresponding Bd update rule Bd^*p*^ leads to uniform initialisation, whereas dB^*p*^ implies temperature initialisation. Table C lists the combinations of update rules and the choices of the individual moving to the vacant site. In the rest of the paper, the following color-coding has been used for the update rules: pink for Bd^*o*^, red for dB^*o*^, green for Bd^*p*^ and orange for dB^*p*^.

To explore other possibilities, we consider the case where the parent moves to the neighboring node and the offspring stays at the original node, see Fig. 1 B. These update rules will be denoted by the superscript *p*. There are biological scenarios that can inspire such a parent moving rule: For example, most lizards and snakes abandon their eggs after laying [43] at specific sites [44–46]. Thus, the rules where the parent moves can be inspired by lizard/snake populations with a fixed number of sites. Movement and reproduction get coupled in birth-death models because there are generally no free spots in a tightly packed populations of fixed size, and new spots open up only after the death events. However, parent moving rules rules are probably more relevant for microbial and somatic cell populations, where parent and offspring cannot necessarily be distinguished, see Fig. 1. In such cases, there is no a priori reason why e.g. a new daughter cell is leaving the mother cell’s site and not vice-versa. There are numerous studies in the mathematical oncology literature that investigate mutation-selection dynamics on regular graphs [47, 48]. In all of these studies, the mutant daughter cell moves to the vacant site, but there is no reason why the daughter cell identical to the mother cell cannot move instead. On regular graphs, the initial placement of the mutant is independent of the choice of the individual moving to vacant site – parent (daughter cell resembling the mother) or offspring (mutant daughter cell). This is not true for non-regular graphs, where the initial placement of the mutant is determined by the choice of individual moving to vacant sites (Fig. 1).

The probability that the initial mutant under dB^*p*^ update arises in node *i* is

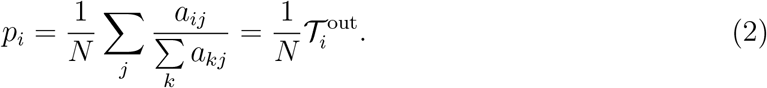

Thus, the mutants for dB^*p*^ updating with parent-type offspring moving to a vacant node are out-temperature initialised. In other words, highly connected nodes are more likely to receive an initial mutant. For unweighted and undirected graphs *𝒯*^in^ = *𝒯*^out^. For weighted graphs, in the definitions of *𝒯*^in^ and *𝒯*^out^, all elements of the adjacency matrix must be replaced with the elements of the weight matrix, **W**. Note that for the complete graph the choice of individual moving to vacant node is irrelevant due to the symmetry of the graph.

In the literature, it has been suggested that for dB updating temperature initialisation does not exist [49, 50]. This is true when the offspring individual moves to a neighbouring node. But when we instead assume that the parent moves, we obtain the temperature initialised dB update. Moreover, Bd^*p*^ updating with the parent moving to the vacant node recovers the uniform mutant initialisation. Table 1 C lists the combination of update rules and the choice of individual moving to the vacant site leading to different mutant initialisation schemes.

## III. SHORT TIME SCALES: FIXATION DYNAMICS

In evolutionary graph theory, the focus is typically on the fixation probability of a mutant. For a given graph, this probability may depend on the node where the mutant first appears.

To make this explicit, we denote by φ_*G,i*_(*f*′, *f*) the fixation probability of a mutant with fitness *f*′ in a wild-type population of fitness *f* on a graph *G* starting from node *i*. As discussed above, the mutant usually does not arise in every node with the same probability.

For a general mutant initialisation scheme *ℐ*, the average fixation probability on graph *G* is

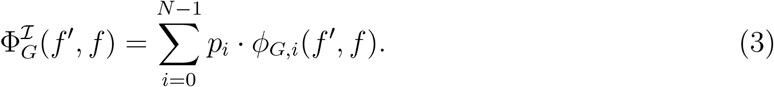

In the following, we study average fixation probabilities on the star graph and on random graphs for different update rules.

### A. The star graph

The star graph has been extensively studied in evolutionary graph theory, as it is highly inhomogeneous but still analytically tractable [51]. The Moran Bd process on the star graph was studied in [13]. Since then, the star graph has become the prime example of an *AoS* for uniform mutant initialisation, which can be solved exactly [52, 53] for various update rules, including some that have not been discussed here like bD and Db [54]. However, the star graph fails to amplify selection if the initial mutant is initialised according to the temperature initialisation scheme, where the central node is more likely to receive the initial mutant [36]. Under temperature initialised Bd updating, the star graph is instead a *SoF* [40], see Fig. 2 A and a *SoS* for uniform initialised dB updating [49], see Fig. 2 B.

**FIG. 2.**
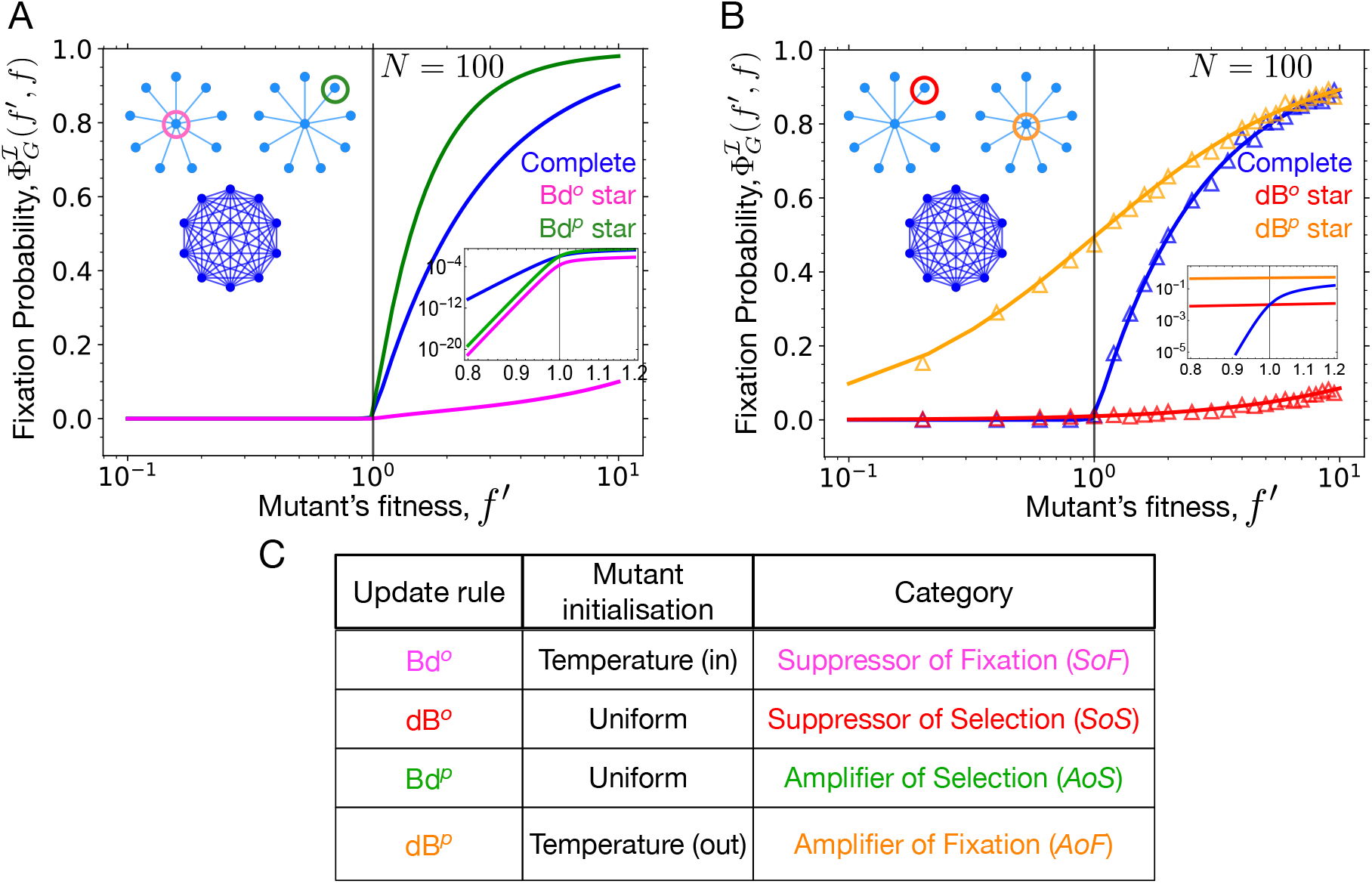
Fixation probability for the star graph under different update rules. A) Under temperature initialised Bd updating (Bd^*o*^), the star graph is a suppressor of fixation whereas, under uniformly initialised Bd updating (Bd^*p*^) the star is an amplifier of selection. B) The uniformly initialised star graph under dB updating (dB^*o*^) is a suppressor of selection. A new category of graphs, amplifiers of fixation, is introduced here. An amplifier of fixation has higher fixation probability for a mutant, regardless of its fitness, than on the complete graph. Under temperature initialised dB updating (dB^*p*^) the star graph is an amplifier of fixation. Specifically, for finite *N* the star graph is a piecewise amplifier of fixation, and only in the limit of infinite population size, it becomes an universal amplifier of fixation. Symbols correspond to dB simulations with 2000 independent runs for each graph (with each run conditioned on fixation). C) Summary of how the choice of update rule affects the fixation dynamics for the star graph.

So far, the star graph was not studied for the temperature initialised dB updating, because this requires the assumption that the parent moves instead of the offspring. For the dB update rule, the fixation probability for a mutant with fitness *f*′ in the background fitness *f* on the complete graph is [15, 33]

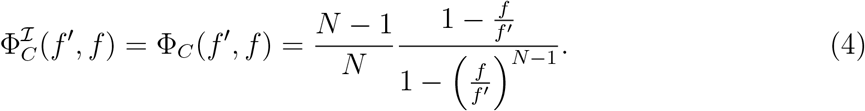

In the limit of large *N* this becomes equal to the Bd fixation probability [15]. Also, because of the symmetry of the complete graph, the fixation probability is independent of the mutant initialisation scheme *ℐ*. This is true for other regular graphs as well. Using the approach of recursive relations [54], in *Appendix* A we derive the fixation probability of a mutant under dB updating on the star graph. When the mutant is initially placed on the center node, its fixation probability is

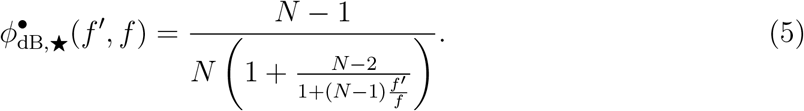

Here the star symbol denotes the star graph, and the filled circle in the superscript of 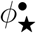 indicates that the initial mutant is at the center node. Similarly, the fixation probability for a mutant initially placed on a leaf node is

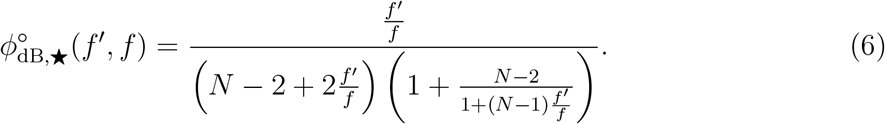

From Eq. 5 and 6, we can compute the temperature initialised fixation probability of a mutant on the star graph under dB updating (equivalent to dB^*p*^) as

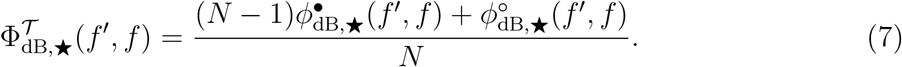

From Eq. 7, we find that for dB^*p*^ updating, the probability to fix advantageous mutants on the star graph is higher than in the well-mixed population, see Fig. 2 B. But the fixation probability for deleterious mutants is also higher than the fixation probability on the complete graph, contrary to the original definition of *AoS* s where the probability to fix deleterious mutant is lower [13]. Therefore, it represents a new category of graphs which we call amplifier of fixation (*AoF*), where the probability to fix a mutant is higher than that of the complete graph, regardless of the mutant fitness value.

The star graph under temperature initialised dB updating dB^*p*^ is a piecewise *AoF* for finite *N*, see Fig. A.1, and only in the limit *N*→ ∞, it is a universal *AoF*, see Fig. 2 B. Intuitively, in a very large population, the initial mutant will most likely appear at the central node. Assuming the mutant does appear at the central node, in the next step an individual is selected with uniform probability to die. Most likely this is a leaf node. In this case, the mutant at the central node replaces the dead individual. This way, the initial mutant can survive in the population with higher probability, regardless of its fitness. Taking the *N* → ∞ limit of the fixation probability profile we find,

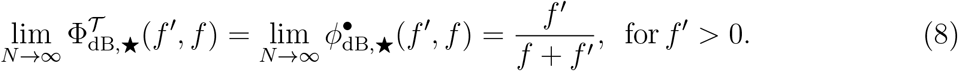

Thus the probability to fix a deleterious mutant (*f*′ *< f*) remains non-zero even in the limit of *N*→ ∞, which contradicts the conventional intuition that deleterious mutants are efficiently purged from large populations. A similar phenomenon was observed in [9, 55] in an explicitly spatial setup. Before we proceed, it is important to note that the choice of individual type moving to vacant sites – parent or offspring – only affects the initial mutant placement on the structure. In the absence of mutations, these two choices are equivalent as both the offsprings resemble the parent type.

### B. Numerical classification of graphs

In the previous section we have analysed the star graph under 4 different updating schemes, Bd^*o*^, dB^*o*^, Bd^*p*^ and dB^*p*^. Depending in the update scheme, the star graph can be a *SoF*, a *SoS*, an *AoS*, or an *AoF*. The classification carries over to other graphs and we implement it as follows: we numerically compute the fixation probability for a mutant with fitness values *f*′ = 0.5, 0.75, 1, 1.25, 1.5, 1.75, 2, 2.25 and 2.5, see *Appendix* B for more details. With wild-type individual fitness *f* = 1, we classify a given connected graph *G* as

- *AoS*, if 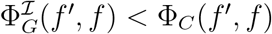 for *f′* = 0.5, 0.75 and 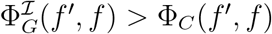 for *f′* ≥ 1.25.
- *SoS*, if 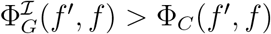 for *f′* = 0.5, 0.75 and 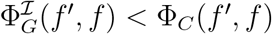 for *f′* ≥ 1.25.
- *SoF*, if 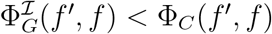 for all *f′*.
- *AoF*, if 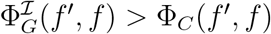 for all *f′*.
- *Piecewise AoF*, if 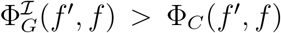 for *f′* ≤ *f* ^∗^ and 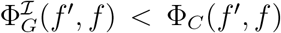 for *f′ > f* ^∗^, where *f* ^∗^ ≥ 1.
- *Isothermal* graph if 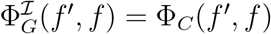 for all pairs of *f′, f*, and every node has the same degree. However, the isothermal graphs have the same fixation probabilities as the complete graph only for Bd updating [35] (see *Appendix* C for further discussion).

For weighted graphs, all isothermal graphs (graphs where every node has equal in-temperature) have the same fixation probabilities under Bd updating as the complete graph. Note that these definitions assume that at neutrality (*f′* = *f*) all the graphs have the fixation probability 1*/N*, as the complete graph which is chosen as the basis for comparison. However, this is true only under uniform mutant initialisation. Under temperature mutant initialisation, the fixation probabilities for graphs at neutrality need not be 1*/N* [56]. Piecewise amplifiers and suppressors are sometimes referred to as transient in the literature [49, 57].

### C. Random graphs

Based on this, we study the fixation probability profiles for random graphs of size 8 for the same update rules, to see to what extent the observations made for the star graph in Sec. III A extend to random graphs. For this purpose, we randomly generated Erdős Rényi graphs [58] for different probabilities of link connection, *p*. Setting *p* = 0 generates fully disconnected graphs whereas *p* = 1 generates the complete graph. As the fixation probability is defined only if the graph is connected, we condition on connected graphs [59, 60]. From Fig. 3 A, we find that, just as the star graph, most random graphs are *SoF* under the temperature initialised Bd process (equivalent to Bd^*o*^). Similarly, most of the random graphs under temperature initialised dB (equivalent to dB^*p*^) are (piecewise) *AoF*, see Fig. 3 B. Therefore, *AoF* and *SoF* are ubiquitous. Under the uniformly initialised Bd process (equivalent to Bd^*p*^) and dB process (equivalent to dB^*o*^), most of the random graphs are *AoS* and *SoS* respectively, see Figs. 3 C,D. The ubiquity of these categories has been shown earlier in [33].

**FIG. 3.**
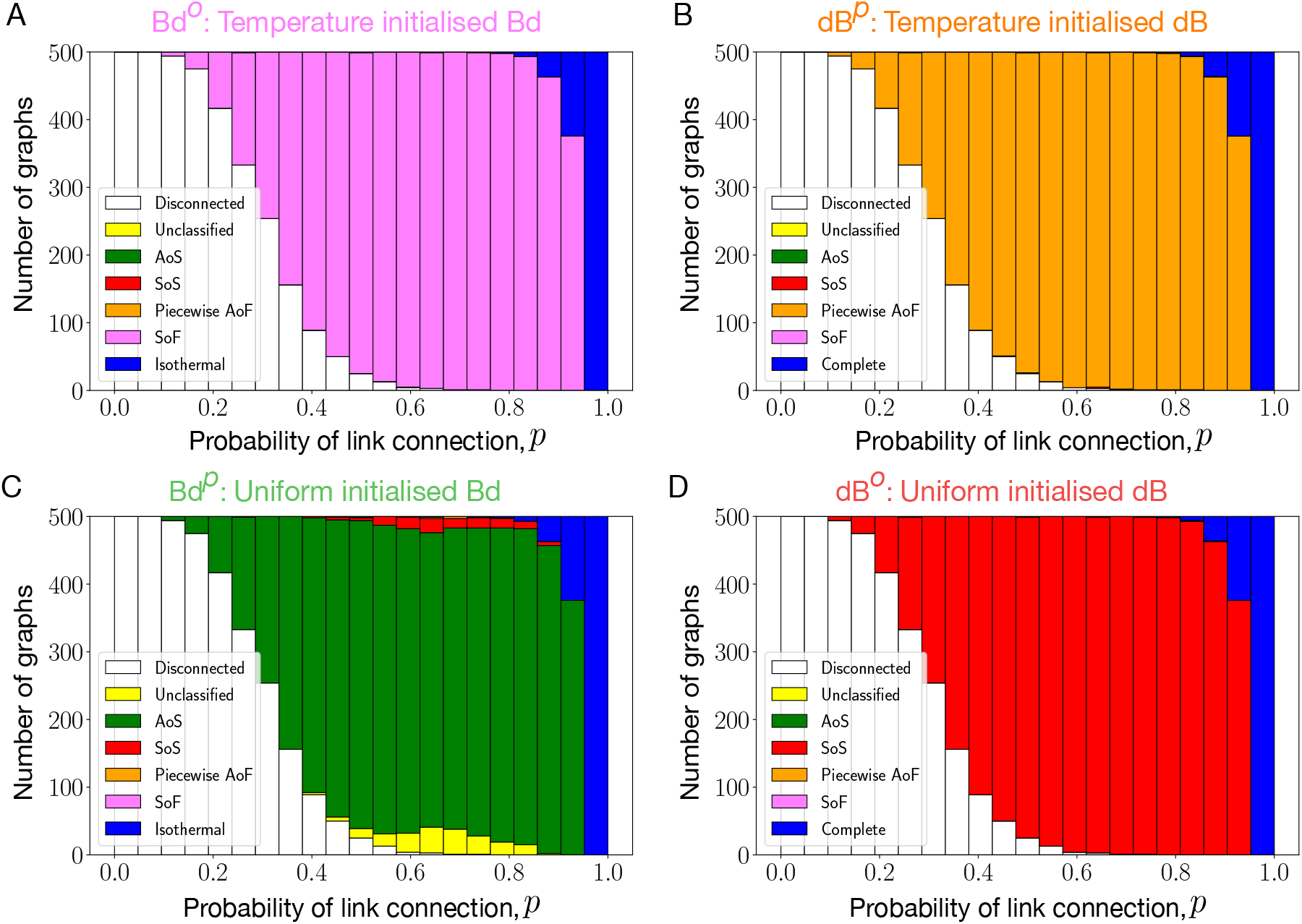
Suppressors and amplifiers of fixation are ubiquitous. We generated Erdős Rényi graphs of size *N* = 8 for several values of the probability of link connection, *p*. Fixation probability profiles for connected graphs are numerically obtained for temperature/uniform initialised Bd and dB updating. Refer to Sec. III B for details on the fitness discretisation. A) Most of the graphs are suppressors of fixation under temperature initialised Bd updating, i.e., most of the graphs have lower fixation probability for a mutant regardless of its fitness than the complete graph, whereas, B) most of the connected random graphs are piecewise amplifiers of fixation under temperature initialised dB updating. These graphs have higher fixation probability than the complete graph for mutants with fitness *f′* ≤ *f* ^∗^, and lower fixation probability for *f′ > f* ^∗^ with *f* ^∗^ ≥ 1. In other words, within our chosen resolution of the fitness scale, the fixation of deleterious mutations is amplified and the fixation of beneficial mutations is suppressed. Similarly, C) most of the connected graphs are amplifiers of selection under uniformly initialised Bd updating, whereas, D) most of the connected random graphs are suppressor of fixation under uniformly initialised dB updating. In Fig. B.1, the fixation probabilities of random graphs with *p* = 0.5 are shown for different update schemes.

## IV. LONG TIME SCALES: MUTATION-SELECTION BALANCE

After studying short-term fixation dynamics in graph-structured populations, we now move our focus to long-term mutation-selection dynamics. We assume that the state space is a bounded fitness interval [*f*_min_, *f*_max_]. During a birth event, with probability 1− *µ* the offspring resembles its parent and has the same fitness, and otherwise it mutates to a new fitness sampled from a mutational fitness distribution *ρ*(*f′, f*) [61]. Here *ρ*(*f′, f*) denotes the probability density of the mutant offspring fitness *f′* given the parental fitness *f*, which is also known as the neighbour fitness distribution [62].

We work in the regime of low mutation rates, *µ*«1 where the population is monomorphic almost all the time, except during a fixation event. All individuals have the same fitness value, which can be used to label the entire population. Specifically, the average time between two successive mutations is large enough so that the initial mutant reaches fixation or goes extinct before the next mutation appears [63]. Any change in the state of the population requires the fixation of a new mutation. Thus, the fixation probability and the mutant initialisation (depending on the details of the update rule) 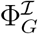 fully determine the long-term mutation-selection dynamics. These dynamics are known by multiple names, e.g., sequential dynamics, periodic selection [25, 64] or origin-fixation dynamics [65].

The origin-fixation dynamics on a population structure *G* is a continuous time Markov chain on the fitness state space governed by the master equation

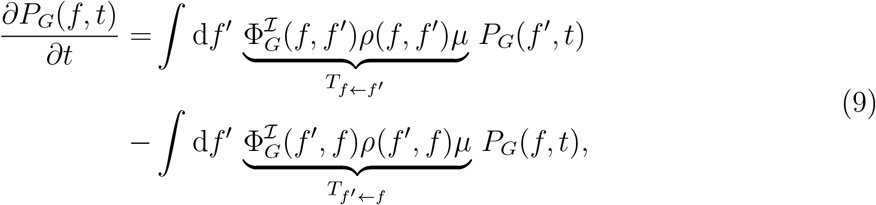

where *P*_*G*_(*f, t*) is the probability density function for the structure *G* to be in between fitness state *f* and *f* + d*f* at time *t*.

At long times a steady-state fitness distribution 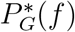 is attained. 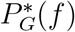 satisfies the stationarity condition

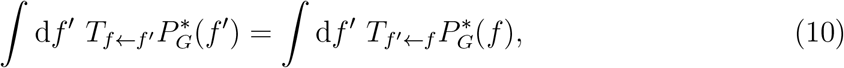

where *T*_*f*←*f′*_ is the transition probability from the fitness state *f′* to *f* and *T*_*f* ←*f*_ is the transition probability from the fitness state *f* to *f′* as defined in the Eq. 9. This condition simplifies considerably if the Markov chain is reversible, which implies the detailed balance relation [66, 67]

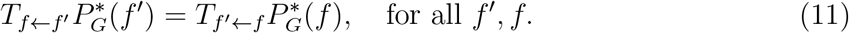

Normalising 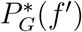, the steady-state solution takes the form,

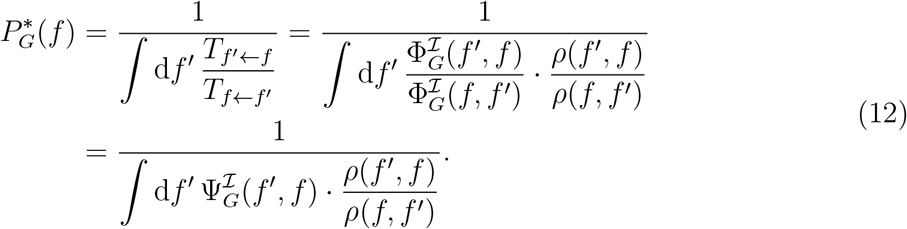

Here, we have introduced the ratio of fixation probabilities 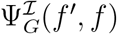. In [68] it has been shown that the origin-fixation dynamics is reversible if and only if 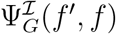 is a power-law, i.e.,

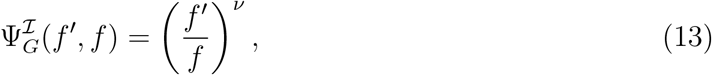

where ν is constant. For most graphs this condition is not satisfied exactly, but it may hold approximately in a range of fitness values and/or for large population sizes, see *Appendix* D for more details. Eq. 12 can then be used as an approximation to the steady state fitness distribution [40].

### A. Complete and regular graphs

For the complete graph under dB updating, 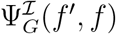 takes the form

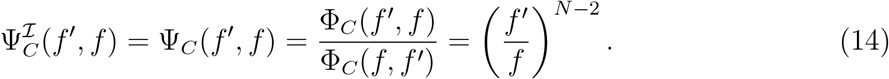

Thus, the Moran dB origin-fixation dynamics on the complete graph is reversible with ν = *N* − 2. The steady-state fitness distribution is therefore given by

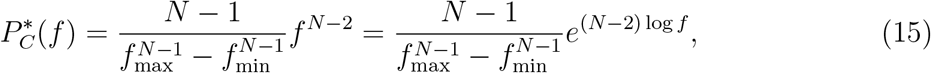

where we assume that the mutant’s fitness is sampled from a uniform distribution, i.e., 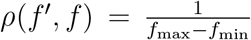. This choice is similar to the House of Cards model [69]. Similarly, for the Moran Bd origin-fixation dynamics, in [68, 70] reversibility is shown to hold for the complete graph with ν = *N*− 1. From now onwards, we use the notation *P*_*G*_ for 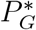.

The steady-state average fitness for the Moran dB origin-fixation dynamics on the complete graph is

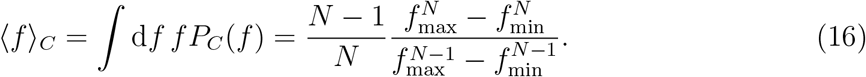

In the limit *N* → ∞,

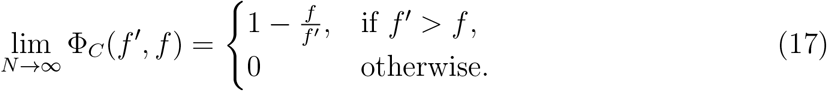

Thus, under this dynamics, an infinitely sized well-mixed population can only move forward on the fitness space with long-term fitness converging to *f*_max_. This can also be seen by performing the *N*→ ∞ on the average steady-state fitness in Eq. 16. The same result holds for the Moran origin-fixation Bd dynamics.

The steady-state fitness distribution in Eq. 15 takes the form of an exponential, Boltzmann-like distribution for −log *f*. This suggests an analogy between statistical mechanics and evolutionary theory that has been pointed out in numerous former studies [70–74]. Under this analogy, a physical system’s energy is equivalent to the negative logarithm of fitness, and the inverse physical temperature is equivalent to the effective population size. Just like high physical temperatures result in strong thermal fluctuations, low effective population sizes lead to highly stochastic population dynamics.

For general (reversible) origin-fixation dynamics, the role of *N* − 2 or *N* −1 for the Moran model on the complete graph is taken over by the exponent ν in Eq. 13. This suggests to define ν as a measure of the effective population size of a general graph (see [12, 75, 76] for definitions of effective population sizes in other contexts). While this definition is strictly applicable only if the long-term origin fixation dynamics is reversible, it is useful also when reversibility holds approximately or asymptotically. We will see below that the definition is consistent with intuition, in the sense that graphs with larger (smaller) effective population sizes display smaller (larger) fluctuations in the evolutionary dynamics. We also studied the long-term dB mutation-selection dynamics for other regular graphs, see *Appendix* C for details. The one-dimensional cycle graph, a *SoF* under dB updating [33, 57], has a lower probability of fixing mutants regardless of the fitness of the mutant compared to the well-mixed population. The cycle graph is worse at fixing beneficial mutations but it is better at preventing the fixation of deleterious mutants. The ratio of the fixation probabilities Ψ_○_ is exactly the same as that for the complete graph, i.e.,

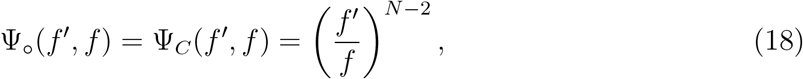

see Eq. C2. In the long run, the lower probability of fixing beneficial mutants for the cycle graph gets compensated by the higher probability of rejecting deleterious mutants. As a result, under Moran dB origin-fixation dynamics, the steady-state statistics for the cycle graph is the same as for the complete graph. Similarly, the two-dimensional grid with periodic boundary conditions, although having a different fixation probability profile [35], approximately attains the same steady-state statistics for the long-term mutation-selection dynamics as the complete graph, see *Appendix* C for more details. Therefore, for long-term dB dynamics we expect other (large) regular graphs to have the same steady-state as the complete graph. For Moran Bd updating, the complete graph, the cycle graph and the two-dimensional lattice (with periodic boundary conditions) have the same fixation probabilities [13], therefore they all have the same steady-state statistics for the Moran Bd origin-fixation dynamics.

### B. Star graph

We now study long-term mutation-selection dynamics on the star graph under Moran Bd^*p*^, Bd^*o*^, dB^*p*^ and dB^*o*^ updating. In the long-term Moran dB^*o*^ origin-fixation dynamics, the complete graph leads to a higher average fitness than the star graph, see Fig. 4 B. This is expected, as the star graph under dB^*o*^ updating is a *SoS*.

**FIG. 4.**
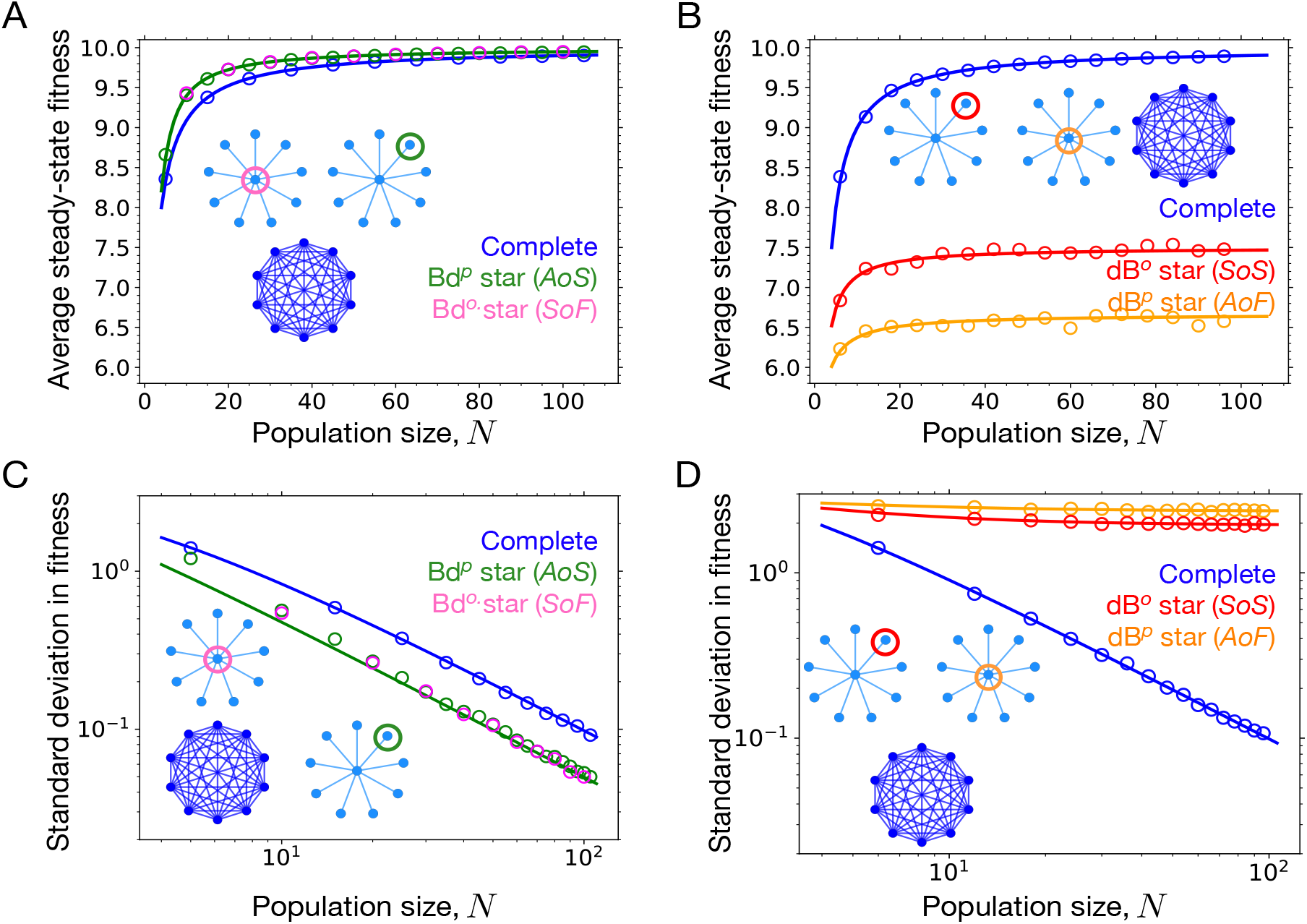
Moran origin-fixation dynamics on the star graph. The figure shows the average and standard deviation of the steady state fitness distribution as a function of population size. Circles represent results obtained from Monte Carlo simulations of the origin-fixation Markov chain, and lines were computed from the approximate expression Eq. 12 for the steady state fitness distribution. A) For Moran Bd^*o*^ origin-fixation dynamics, the star graph, despite being a suppressor of fixation, attains not only higher average fitness than the complete graph, but identical fitness as the star graph under Bd^*p*^ dynamics, where it is an amplifier of selection. This happens because the Bd^*o*^ star compensates for its inability to fix beneficial mutants by rejecting deleterious mutants efficiently. B) For dB^*p*^ dynamics, the star graph, being an amplifier of fixation, not only attains lower steady-state average fitness than the well-mixed population, but also lower than the star graph subjected to dB^*o*^ update, where it is a suppressor of selection. This happens because the dB^*p*^ star is worse in rejecting deleterious mutations than the dB^*o*^ star. Therefore, being good at fixing beneficial mutants is not sufficient to attain higher fitness in the long-term evolution. C) The star graph under Bd long-term dynamics not only attains higher average fitness but also lower fluctuations in the steady-state than the well-mixed population. This can be understood by the higher effective population size of the star graph. D) Compared to the average fitness order under dB dynamics in panel B, the order for the standard deviation in fitness is reversed: dB^*p*^-star, Bd^*o*^-star and the complete graph. Moreover, the standard deviation for the star graphs under dB long-term dynamics saturates to finite values for large *N*, as their effective population sizes are independent of *N*. Parameters: *f*_min_ = 1, *f*_max_ = 10 and the number of independent runs for Monte Carlo simulations is 2000.

For the Moran dB^*p*^ origin-fixation dynamics, the star has a higher probability to fix mutants than the complete graph. However, in the long-term dynamics the complete graph leads to a higher average fitness than the star graph. Moreover, the star graph with dB^*o*^ updating attains higher fitness than the star graph subjected to dB^*p*^ updating. This happens because the star graph under dB^*p*^ updating has a higher probability to fix deleterious mutants. As the population gets closer to the fitness peak (here *f*_max_), the probability for the mutations to have deleterious fitness effects also increases. Consequently, the fate of deleterious mutants has a strong influence on the steady-state fitness of the population.

To understand this further, we study the large *N* behaviour of the steady-state for the various Moran origin-fixation dynamics on the star graph. In the limit of large *N* for Moran dB^*p*^ updating, we find from Eq. 8 that

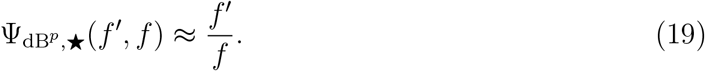

From Eq. 19, we find that the effective population size in the large *N* limit for the star graph under dB^*p*^ updating is ν = 1. The corresponding steady-state fitness distribution in the large *N* limit is

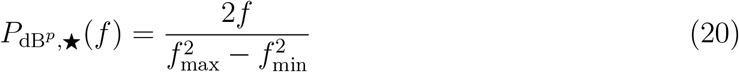

and the steady-state average fitness is

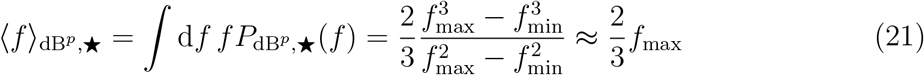

for *f*_max_ ≫ *f*_min_. Similarly, the large *N* limit for the star graph under dB^*o*^ updating gives (see *Appendix* A for details)

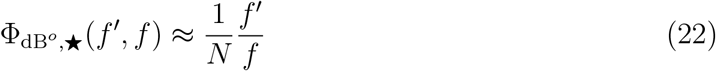

and therefore

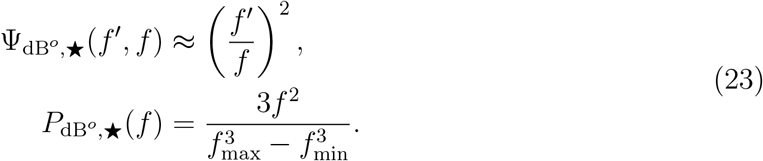

In the limit of large *N*, the effective population of the star graph subjected to dB^*o*^ dynamics is twice that obtained for dB^*p*^ dynamics. As a consequence, the dB^*o*^ star graph attains higher average fitness in the steady-state,

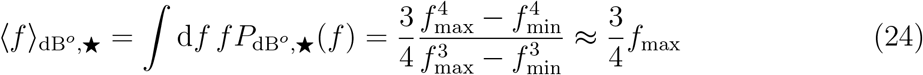

for *f*_max_ ≫*f*_min_. Although 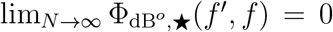 and 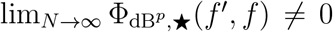, we find 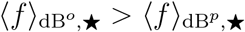 (but note that, because of the lower fixation probability, the higher steady state fitness of the dB^*o*^ star is attained after a longer time).

Performing a similar analysis for Birth-death updating, we find that the star graph under Bd^*p*^ and Bd^*o*^ updating satisfies

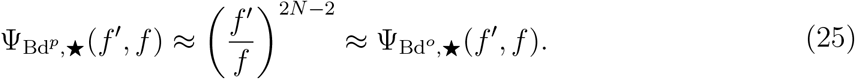

The first approximation follows from [52] and the second approximation follows from [40]. What this means is that, although under Moran Bd^*p*^ and Bd^*o*^ updating the star has quite different fixation probability profiles, see Fig. 2 A, in the long term to a good approximation they display identical steady-state statistics, because in both cases the star graph has the same effective population size of 2*N*. This also means that the star graph under long-term Moran Bd dynamics attains higher average fitness in the steady-state than the well-mixed population. Specifically, the steady-state fitness distribution is

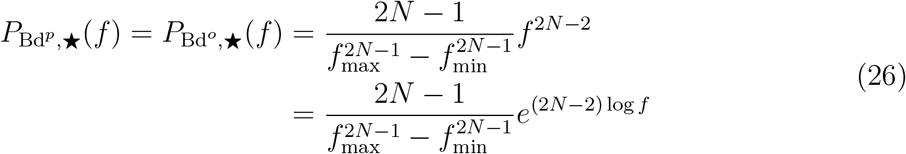

and the average steady-state fitness is

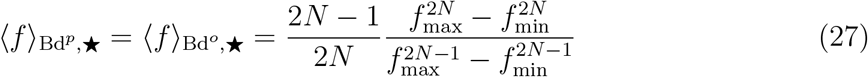

with

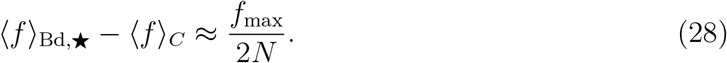

The star graph under Bd^*p*^ updating, an *AoS* (Fig. 2 A), expectedly attains higher fitness than the well-mixed population because it is better in fixing beneficial mutations and preventing the fixation of deleterious mutations. However, the star graph under Bd^*o*^ updating, a *SoF*, also attains a higher steady-state average fitness than the complete graph (identical to the one under Bd^*p*^ updating) because it is much better at rejecting deleterious mutants, which compensates for its lower probability to fix beneficial mutations [40].

The effective population size also affects the fluctuations in the steady-state. Because the effective population size of the star graph under dB updating is independent of *N*, the standard deviation in fitness does not change at large *N*. The *AoF* star experiences higher fluctuations than the *SoF* star because of its lower effective population size, see Fig. 4 D. Under Bd updating, the effective population size of the star graph is twice the effective population size of the complete graph. Therefore, the star experiences lower fluctuations than the complete graph and the standard deviation decreases with increasing *N*, see Fig. 4 C. For more details, see *Appendix* E.

### C. Random graphs

For long-term evolution on the star graph, the deleterious mutant regime can substantially affect the fate of the dynamics. Does this effect extend to other graphs? Can we expect the *SoF* that we found in Fig. 3 A to have higher long-term fitness than the well-mixed population? The answer in not obvious. Similarly, what can we say about the long-term fitness fate of the *AoF* found in Fig. 3 B? Do all of them attain lower steady-state average fitness than the complete graph, just like the star graph under dB^*p*^ updating? We explore these questions next. We move forward by discretising the fitness space and study the Moran Bd^*p*^, Bd^*o*^, dB^*p*^ and dB^*o*^ origin-fixation dynamics for several random graphs. Steady-state statistics of these graphs are obtained by solving the respective Markov chains numerically, see *Appendix* F for details.

Computing the steady-state average fitness for all connected random graphs, in Fig. 5 A we find that for long-term Bd^*o*^ dynamics, almost all *SoF* attain higher steady-state average fitness than the complete graph, whereas in Fig. 5 B for long-term dB^*p*^ dynamics, all the piecewise *AoF* attain lower steady-state average fitness. Interestingly, for the case of Bd^*p*^ dynamics where most of the connected graphs are *AoS*, the graphs attain quite similar average fitness as they do when subjected to the Bd^*o*^ dynamics, see Fig. 5 C. In Fig. 4 A, we have seen that the star graph attains the same fitness for the two kinds of Bd long-term dynamics. Now we confirm this for all other graphs. Expectedly, for the long-term dB^*o*^ dynamics where most of the random connected graphs are *SoS*, the majority of random graphs attain lower average fitness, see Fig. 5 D.

**FIG. 5.**
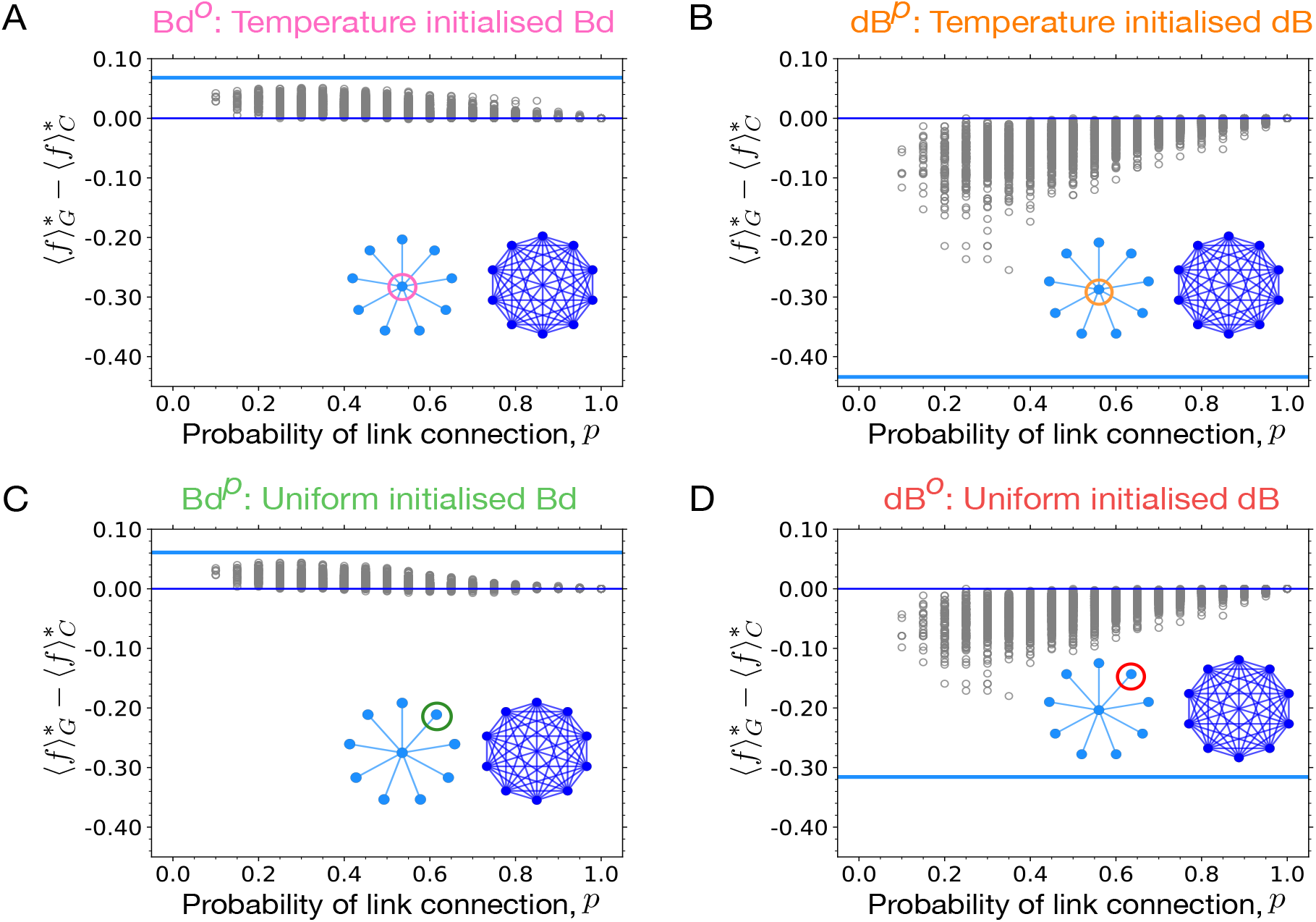
Suppressors of fixation attain higher long-term fitness whereas amplifiers of fixation attain lower fitness. Numerical solutions of the Markov chain on the fitness space for several random graphs of size 8 are presented. The same fitness discretisation is chosen as in Fig. 3. The magenta horizontal lines represent the steady-state fitness obtained on the star graph relative to the complete graph (blue horizontal line) under different dynamics. A) Most of the random graphs are suppressors of fixation under dB^*o*^ updating, yet they attain higher fitness than the well-mixed population. B) Similarly, most of the random graphs are (piecewise) amplifiers of fixation under dB^*p*^ updating, yet they attain lower steady-state average fitness. C), D) Expectedly, amplifiers of selection attain higher fitness for Bd^*o*^ dynamics and suppressors of selection attain lower fitness for dB^*p*^ dynamics. Thus, the deleterious mutant regime is important for generic graph structures when subjected to long-term dynamics.

It is difficult to find general conditions under which a structured population has higher average steady state fitness than the complete graph. In *Appendix* G, we derive a sufficient condition for this. We use this condition to determine the ordering of the average steady state fitnesses for the structures reported in Fig. 4 B and show that they are consistent with the numerical results.

## V. DISCUSSION

Most of the initial research in evolutionary graph theory has focused on the Moran Birth-death (Bd) update with uniform initialisation [13, 26, 52]. The uniform initialisation is typically justified by considering spontaneous mutations during an individual’s lifetime [77]. However, when mutations occur during reproduction instead [36], justifying the use of uniform initialised Bd updating becomes more challenging. In that scenario, temperature initialisation is a more natural choice. Our findings demonstrate that the uniform initialisation in the Bd update naturally arises when parent-type offspring move to vacant nodes (Bd^*p*^) rather than mutant offspring individuals (Bd^*o*^). Furthermore, by necessitating parent-type individuals to move instead of mutant offspring individuals, we have uncovered the existence of temperature-initialised dB updating (dB^*p*^), an update scheme previously considered non-existent. In conclusion, we emphasise that a mutant initialisation scheme is an outcome of an update rule and the mode of mutations, and need not be specified on top of it. An update rule should be sufficient to generate the fixation dynamics on a graph.

Moran dB^*p*^ updating introduces a new category of graphs known as *AoF* (amplifiers of fixation), where the fixation probability is higher regardless of the fitness values compared to the complete graph. The star graph under dB^*p*^ updating is an *AoF* with non-zero probability to fix deleterious mutants, even in the limit of infinite population size. For all other previously known graphs, such as *AoS, SoS, SoF*, the complete graph, the probability of fixing any deleterious mutant goes to zero for large population sizes. Consequently, the deleterious mutant regime has not been extensively explored in the literature. The discovery of *AoF* underscores the need to consider the deleterious mutant regime in graph classification, and its importance in the computation of fixation probabilities and time.

The results derived from the analysis of the star graph for different update rules also extends to Erdős-Rényi random graphs. Specifically, we observe that the majority of small random graphs are *SoF* under Bd^*o*^ updating, and piecewise *AoF* under dB^*p*^ updating. This finding closely resembles the result of Ref. [33], where most random graphs were identified as *AoS* under Bd^*p*^ updating and *SoS* under dB^*o*^ updating. Consequently, it is not only the order of birth and death events, but also the choice of the individual moving to vacant sites significantly impacts the results at the short-term fixation time scale. Earlier work in evolutionary graph theory has focused on designing strong amplifiers of selection – structures with a high probability of fixing beneficial mutants [13, 29, 78, 79]. Our work offers new research directions where structures can be designed to obtain desirable fixation profiles, both for the beneficial and deleterious mutant regime. Different update rules allow to manipulate the fixation probabilities of beneficial and deleterious mutations independently.

The choice of moving individuals also affects the long-term Moran origin-fixation dynamics. The star graph, a *SoF* under Bd^*o*^ updating, despite having lower probability to fix advantageous mutants attains higher fitness in the long-term dynamics than the well-mixed population. Similarly under dB^*p*^ updating, the star graph despite being an *AoF* with higher probability of fixing beneficial mutants attains lower fitness. In the former case, the star graph is better in rejecting deleterious mutants, compensating for its lower probability to fix beneficial mutations. In the later scenario, the star graph is not good is preventing the fixation of disadvantageous mutants despite being better at fixing beneficial mutations.

More concretely, the effective population sizes of the star graph for different updating explain the corresponding steady-states and the contribution coming from the deleterious mutant regime. The *SoF* star graph has a higher effective population size than the *AoF* star graph. Additionally, the results obtained for the long-term evolution on the star graph also extend to random graphs. Under Bd^*o*^ updating, most of the random graphs, despite being *SoF*, attain higher fitness than the isothermal graphs. Whereas, under dB^*p*^ updating, most of the random graphs despite being piecewise *AoF* attain lower fitness than the isothermal graphs.

To summarise, care should be taken before speculating about the fate of long-term evolution on spatial structures based on the short-term fixation dynamics. For a population adapting on a fitness landscape, the outcome depends on two factors. First, how effective the population is in stepping forward, and second, how good it is in not falling backward. The effect of deleterious mutant regime can also be seen in the transient phase of evolutionary dynamics [61]. The likelihood for the average fitness trajectory to be non-monotonic increases with the probability to accept deleterious mutations, see *Appendix* H for more details. Overall, the deleterious mutant regime is important when it comes to long-term evolution on spatial structures, something that is often ignored in adaptive evolution theories with well-mixed populations. As the present work has focused on long-term evolution in the regime of low mutation rates and origin-fixation dynamics, the role of deleterious mutations for structured populations subject to a large supply of mutations should be addressed in future research.

Experiments with microbial populations have begun to systematically compare evolution in well-mixed and structured environments [10, 11, 80], and the weakened selection against deleterious mutations predicted by theory has been confirmed experimentally [8]. Very recently, the first evolution experiments designed to test predictions of evolutionary graph theory have been performed [81], and it is only a matter of time before further studies are conducted along these lines [82]. The quantitative description of such experiments necessitates the extension of evolutionary graph theory to the structured metapopulation level [34, 83–87]. While our work focuses on the setting of one-node-one-individual, it is expected that the results obtained here should be transferable to metapopulations. Indeed, the analysis in this paper can be adapted for the study metapopulations, and we have identified amplifiers and suppressors of fixation among the metastar structures presented in [86], see *Appendix* I for more details. In this way, the results presented in this paper gain empirical relevance and can be tested in experimental settings [82]. Examining the role of the deleterious mutant regime for structured metapopulations is clearly an important future direction.

## ACKNOWLEDGMENTS

We are thankful to Alex McAvoy, Helen Alexander, Yuriy Pichugin, Joshua Plotkin and Arjan De Visser for helpful discussions. This work was supported by the Max Planck Society and by DFG within CRC 1310 “Predictability in Evolution”.

## Appendix A

### Exact formula for the fixation probability of a mutant on the star graph under dB and Bd updating

The state of the population can be described by (•*/*○, *i*) where the first index indicates if the central node is occupied by a mutant (•) or not (○) and the second index gives the number of mutants in the leaf nodes. Let us denote 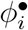 as the fixation probability of the mutant type when started with *i* mutant individuals in the leaf nodes and a mutant at the central node. Similarly, 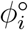 is the fixation probability of the mutant type when started with *i* mutant individuals in the leaf nodes with the central node occupied by a wild-type individual. With *n* number of leaves, 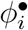 and 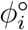 satisfy the following recursion relations [52],

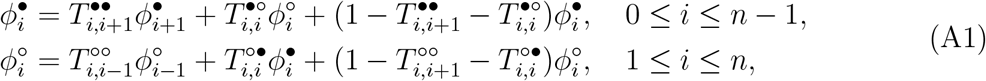

where

− 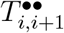 is the probability to transition from the state (•, *i*) to the state (•, *i* + 1).
− 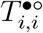 is the probability to transition from the state (•, *i*) to the state (○, *i*).
− 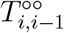 is the probability to transition from the state (○, *i*) to the state (○, *i* − 1).
− 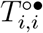 is the probability to transition from the state (○, *i*) to the state (•, *i*).

On rearranging the recursion relations we get,

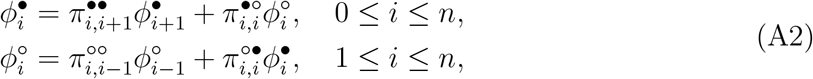

where,

− 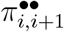 is the conditional probability to transition from the state (•, *i*) to the state (•, *i* + 1) given that the number of mutants changes.

− 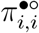 is the conditional probability to transition from the state (•, *i*) to the state (○, *i*) given that the number of mutants changes.

− 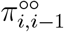 is the conditional probability to transition from the state (○, *i*) to the state (○, *i*− 1) given that the number of mutants changes.

− 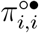 is the conditional probability to transition from the state (, *i*) to the state (, *i*) given that the number of mutants changes.

The conditional probabilities are given by,

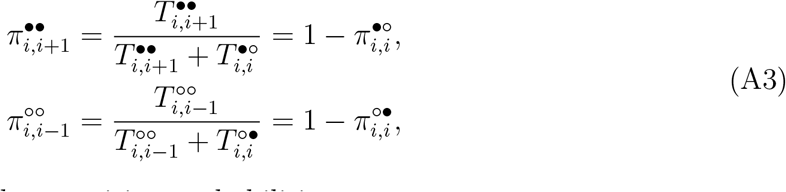

For the Moran dB updating the transition probabilities are,

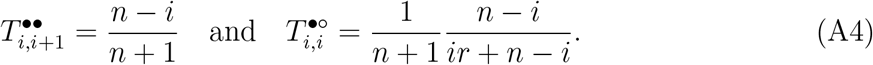

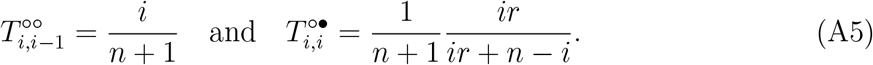

Consequently the conditional probabilities are,

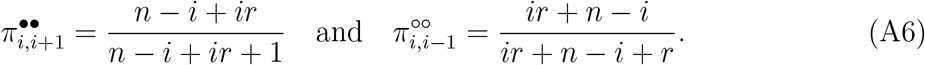

From the ref. [54], we know that the probability of fixation of a mutant appearing at the center node is

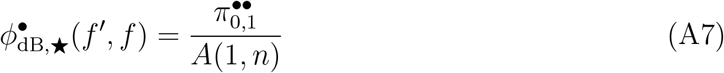

where,

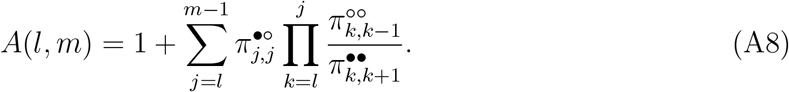

After substituting for the conditional probabilities, we get

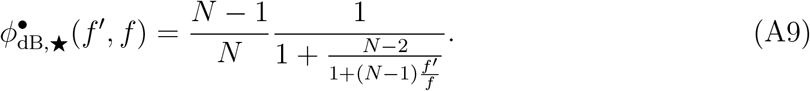

Similarly, we find

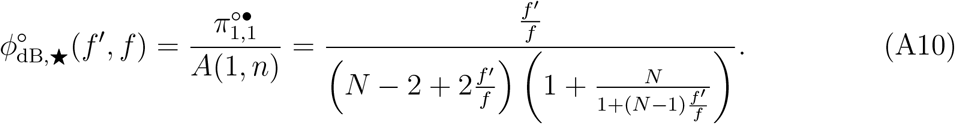

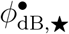 and 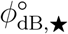 are used to compute uniform and temperature initialised dB fixation probabilities for the star graph.

The uniform (when offspring move to vacant node) and temperature (when parent move to vacant node) initialised fixation probability under Moran dB updating are,

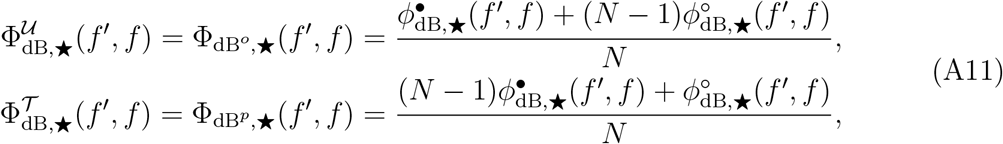

From Fig. 2B (main text), the uniform initialised star graph is a suppressor of selection [49, 57] whereas the temperature initialised star graph is an amplifier of fixation.

Similarly, for Bd updating,

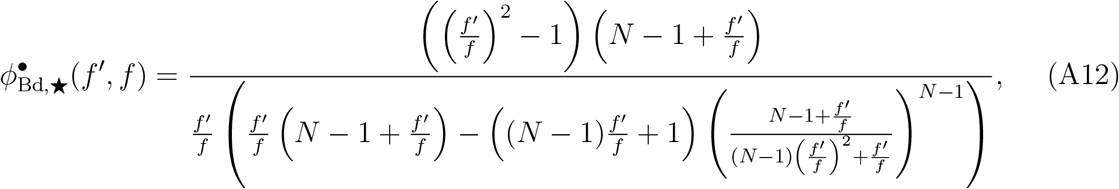

and the fixation probability for a mutant initially placed on a leaf node is,

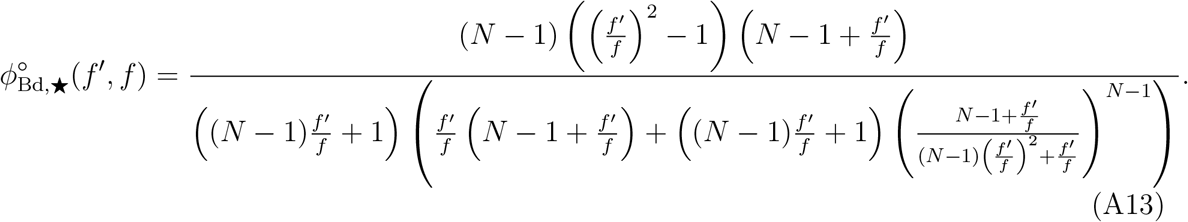

Eqs. A12, A13 are used to compute uniform and temperature initialised fixation probabilities for the star graph under Bd updating.

**FIG. A.1.**
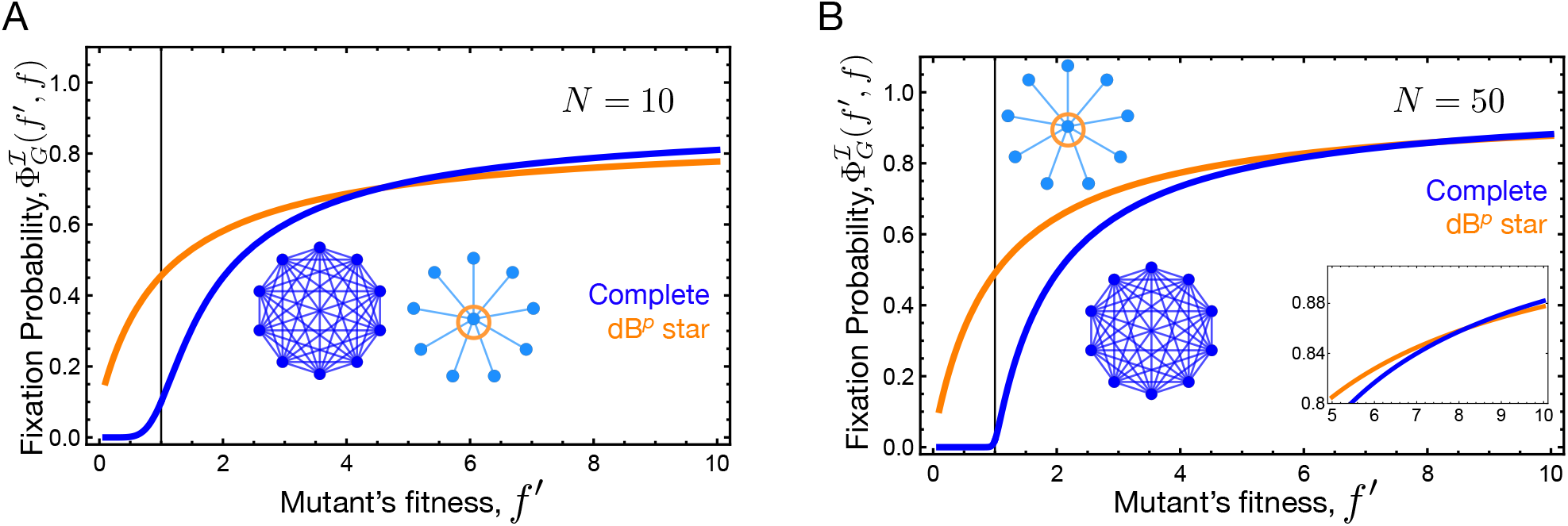
The star graph, a piecewise amplifier of fixation for finite *N*. The star graph under dB^*p*^ updating is a piecewise amplifier of fixation for finite population size *N*. That is for finite *N*, the probability of fixation of a mutant with fitness *f*′ on the star graph is higher than the complete graph for *f*′ ≤ *f* ^∗^ and lower for *f*′ *> f* ^∗^. In the figure we can see that the value of threshold fitness *f* ^∗^ increases on increasing the population size. Only in the limit of infinite population size, the star graph becomes a true amplifier of fixation for dB^*p*^ updating.

## Appendix B

### Matrix approach to compute fixation probability on a random graph

The matrix method solves the Markov chain for the fixation dynamics on a graph. This method is generally used to compute the fixation probability for an arbitrary connected graph [88]. The primary reference for this section is [89]. With states being the configurations of mutant and wild-type, the transition matrix **M**_*s×s*_ is defined as

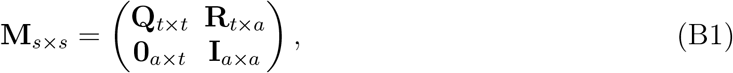

where *s* denotes the total number of states and is equal to *t* + *a*, with *t* being the number of transient states and *a* being the number of absorbing states. **Q** is the transition probability matrix corresponding to the transitions among the transient states, while **R** represents transitions from the transient to the absorbing states. Since by definition there is no jump possible from an absorbing to the transient sector, the lower left matrix is a zero matrix. By similar reasoning, the lower right matrix is an identity matrix.

We denote the fundamental matrix as **F**. It is equal to 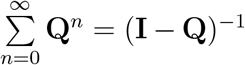. The fixation probability to the absorbing state *j* when started in a transient state *i* is given by the relation,

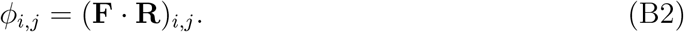

Notice that the indices *i, j* etc., do not represent the number of mutants but the configurations themselves. The second index of the subscript can represent two absorbing states, every individual being the wild-type or the mutant type. It represents the position of the node where the initial mutant appears.

The fixation probability of a mutant to state *j* on a graph *G* with mutant initialization *ℐ* is equal to,

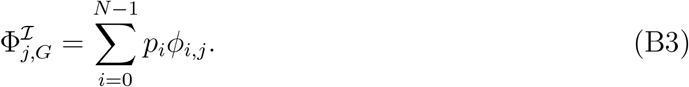

The index *i* in the equation above corresponds to the states where the initial mutant appears at different positions on the graph.

#### a. Transition matrix for the Bd process

For a network of size *N*, we have the transition matrices of dimensions 2^*N*^*×* 2^*N*^. One can decrease the size of these matrices by considering the symmetries of the graph. This has been done in [90] where all the undirected connected networks of size four were considered. After considering the symmetries, the size of the transition matrices for the complete, diamond, and ring graph were reduced from 16 (2^4^) to 5, 9, and 5, respectively. It is not straightforward to account for these symmetries for larger networks; thus, we work with maximum-size transition matrices. We compute the transition matrix for the Moran Bd dynamics with a focus on undirected graphs and unweighted having symmetric adjacency matrices, **A**. We work only with connected graphs as the fundamental matrix **F** becomes singular for the disconnected graphs.

The matrix element **M**_*ij*_ with *i* ≠*j*, the probability of going from state *i* to state *j* with *i* ≠ *j* is given as,

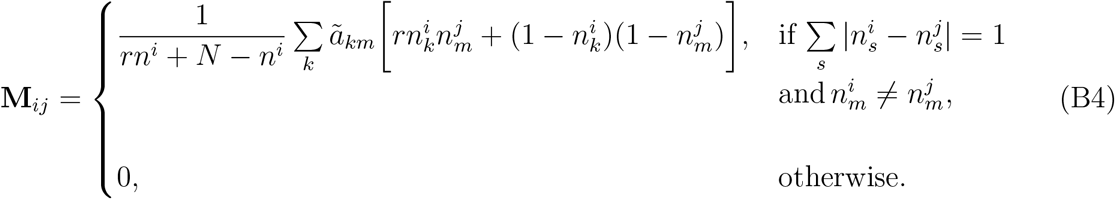

where every node of the configuration *i* can either take a value 0 (for wild-type) or 1 (for mutant). The number of mutants in state *i* is given by 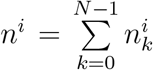. Here, we have also introduced 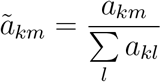. The diagonal elements **M**_*ii*_ are equal to 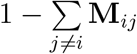.

#### b. Transition matrix for dB process

Like the Bd process, here we write down the transition matrix for the dB process. The matrix element **M**_*ij*_ with *i ≠ j* is given as :

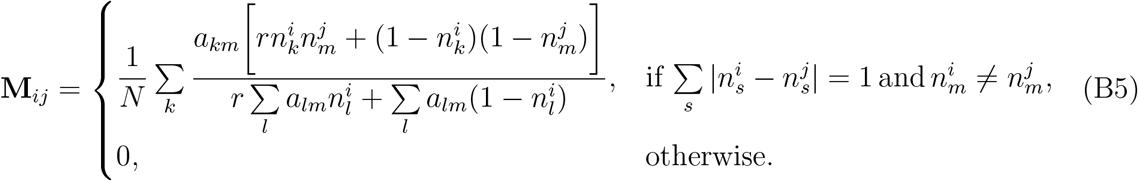

## Appendix C

### Long-term evolution on regular graphs

Under the Moran dB updating, regular graphs have different fixation probability profiles compared to the complete graph [35]. It is in contrast to the Moran Birth-death (Bd) rule, where according to the isothermal theorem [13, 15] all the regular graphs have the same fixation probability profiles as the well-mixed population. For example, under Moran dB updating, the cycle graph has the fixation probability [33],

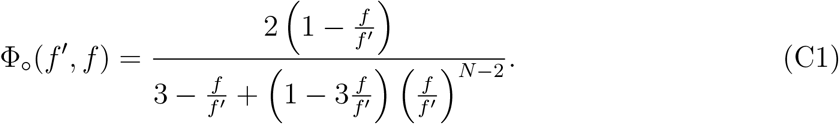

Under the Moran dB updating, the cycle graph is a suppressor of fixation [33], because Φ_○_(*f′, f*) *<* Φ_*C*_(*f′, f*) for all pairs of *f′* and *f* (ignoring the neutral point *f′* = *f*, where 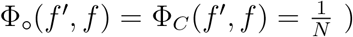), see Fig. C.1 B. The ratio of fixation probabilities entering the steady-state detailed balance solution, however, is the same as complete graph,

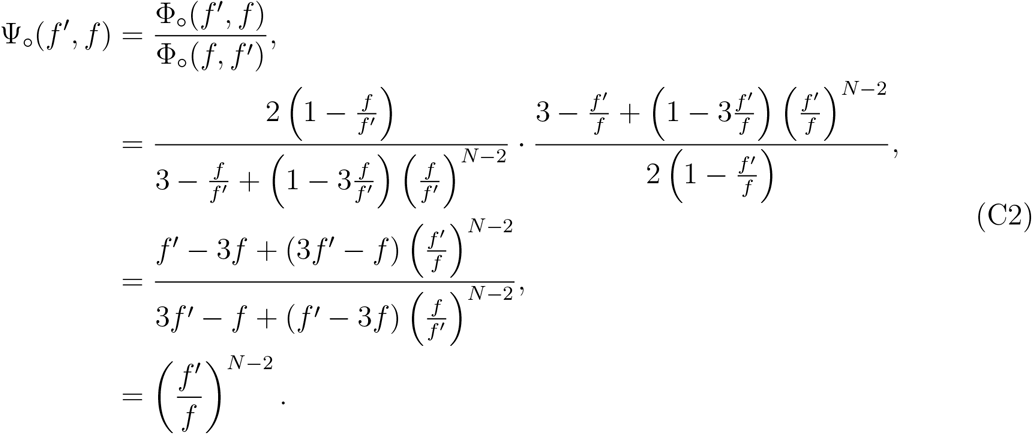

**FIG. B.1.**
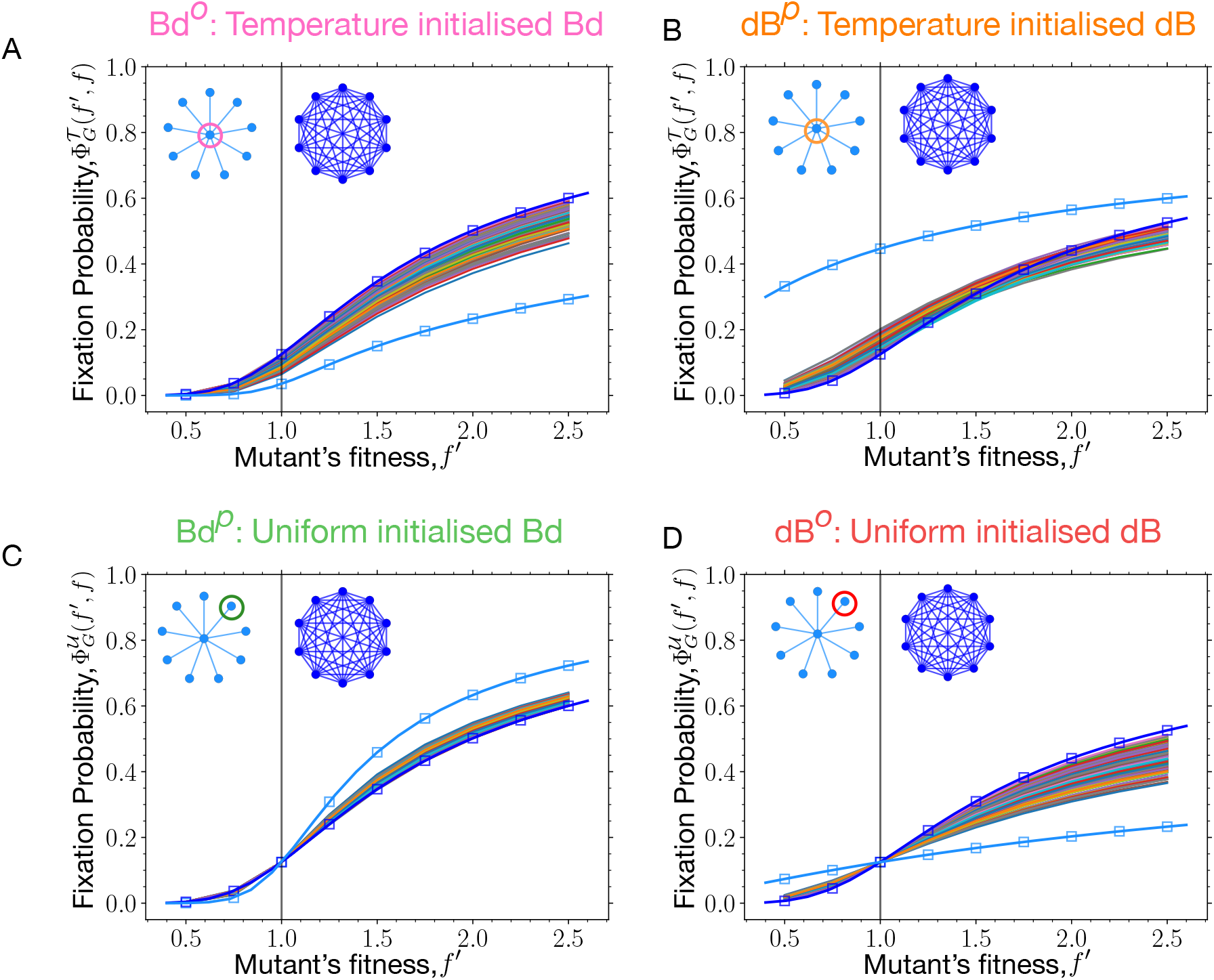
Fixation probability profiles of random graphs. The relative fitness values of the mutant are chosen to be, 0.5, 0.75, 1, 1.25, 1.5, 1.75, 2, 2.25 and 2.5 with wild-type fitness being 1. The fixation probability profiles under different update rules for connected random graphs with the probability of link connection, *p* = 0.5 are shown. The fixation profiles for the complete graph (in blue) and the star graph (in magenta) in both numerical (square markers) and analytical (solid lines) form are also shown. A) Most of the random graphs are suppressors of fixation under temperature initialised Bd updating. B) Most graphs are piecewise amplifiers of fixation under dB^*p*^ updating i.e. the graphs have higher fixation probability for a mutant with fitness below a certain value, *f* ^∗^, and lower fixation probability beyond *f* ^∗^. We observe that beyond *f′* ≈ 1.5, the fixation probabilities become lower than the fixation probability on the well-mixed population. C) Most of the graphs are amplifiers of selection under (Bd^*p*^) uniform initialised *Bd* updating, and D) suppressors of selection under (dB^*o*^) uniform initialised dB updating.

Because Ψ_○_(*f′, f*) is a power law, the Moran dB origin-fixation dynamics on the cycle graph is reversible. Moreover,

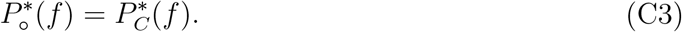

Consequently, 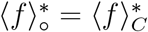.

What did we just learn? We learnt that the cycle graph despite being a suppressor of fixation, attains the same average fitness in the mutation-selection balance as the complete graph, see Fig. C.1 D. In the limit of *N* → ∞,

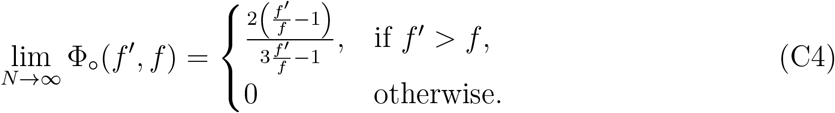

Now,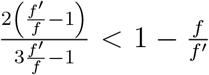 for all *f′ > f*, therefore, in the limit of very large population sizes, the cycle graph is less likely to fix beneficial mutants than the complete graph, see Fig. C.1 A. Yet the cycle and the complete graph attain the same steady-state average fitness for all population sizes. It happens because the cycle graph is better at rejecting deleterious mutants than the complete graph. The ability of the cycle graph to prevent the fixation of disadvantageous mutants compensates for its lower probability to fix beneficial mutants in a way that Ψ_○_(*f′, f*) becomes equal to Ψ_*C*_(*f′, f*).

We also explore the steady-state statistics of the long-term dB mutation-selection dynamics on the two dimensional lattice with periodic boundary conditions. With each node having *k* neighbors, the fixation probability of a mutant to fix on the 2d lattice is [35],

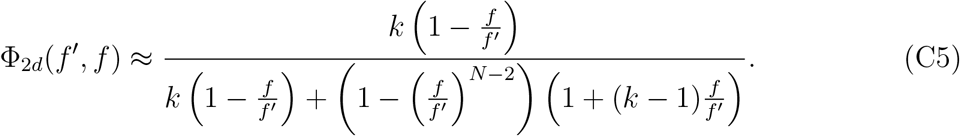

We focus on the case of *k* = 4. The ratio of fixation probabilities takes the form,

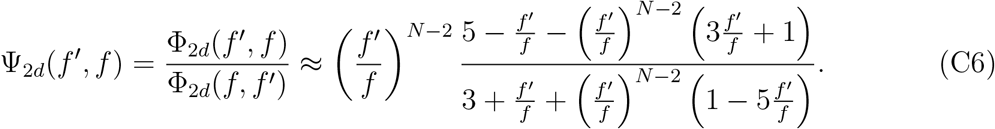

From Fig. C.1 A and C, it is clear that the fixation probability profile of the 2d lattice is different from the well-mixed population, but the graph category to which the 2d lattice belongs is not clear. This could be due to the approximation made in Ref. [35] to compute Eq. C5. As a result, for small *N*, the 2d lattice attains different (lower) steady-state average fitness in the mutation-selection balance than the complete and cycle graph. However, with an increase in population size, the steady-state average fitness attained by the 2d lattice asymptotes to the one attained by the well-mixed population, see Fig. C.1 D. This can be understood by performing the large *N* expansion on Ψ_2*d*_(*f′, f*) yielding,

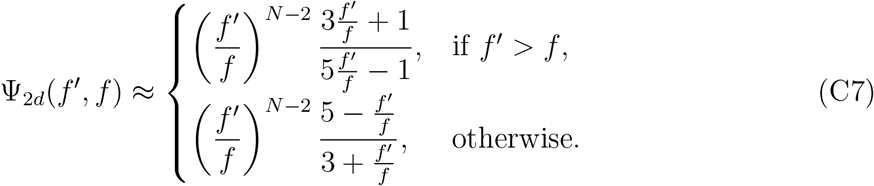

For large population sizes, 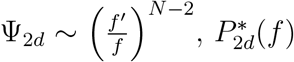 and therefore, 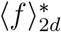 take the same limit as the complete graph. In the argument made, we have assumed reversibility to hold. In the subsequent section, we justify the assumption.

From the above two case studies, we have learnt that although the complete graph, the cycle graph, and the 2d lattice behave differently at fixation time scales under dB updating [35], their steady-state statistics for the long-term Moran dB origin-fixation dynamics are identical.

## Appendix D

### Reversibility

The primary reference for this section is [68]. Using the Kolmogorov criterion [91], the neutral Moran origin-fixation dynamics turns out to be reversible. With this, the origin-fixation dynamics under selection is reversible if 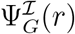 is a power law,

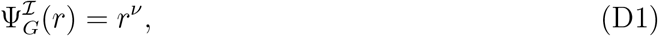

where *r* is used as a shorthand for 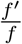, and ν is given by the relation,

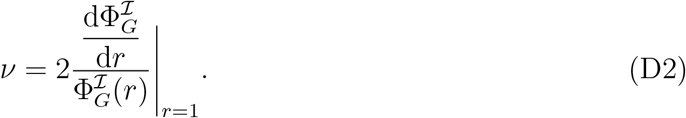

In general, 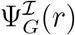 need not be a power law. Therefore, to check the scope for reversibility a logarithmic expansion is performed on 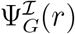,

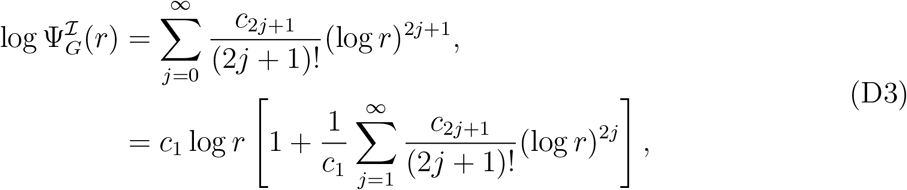

where,

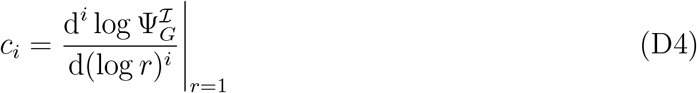

and *c*_1_ = ν. This can be seen as following,

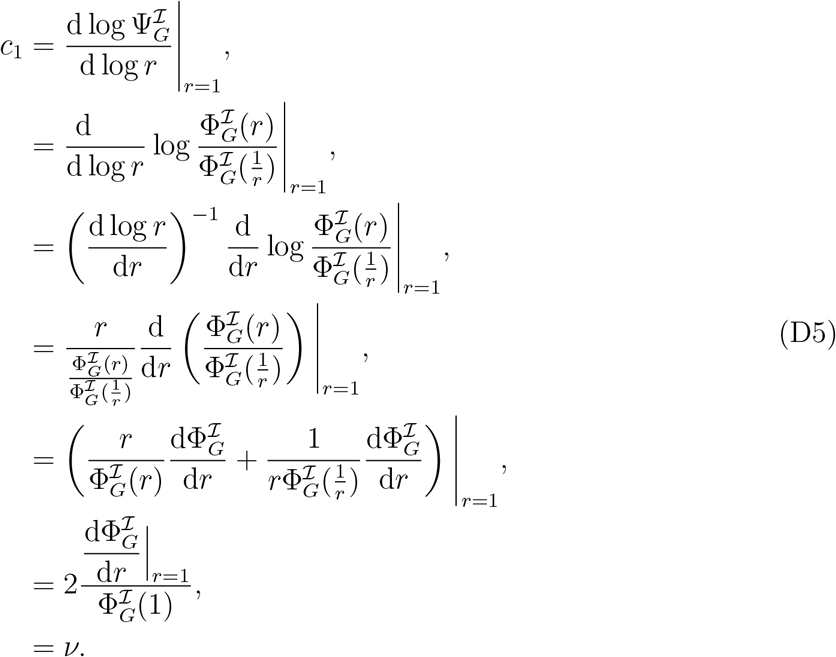

**FIG. C.1.**
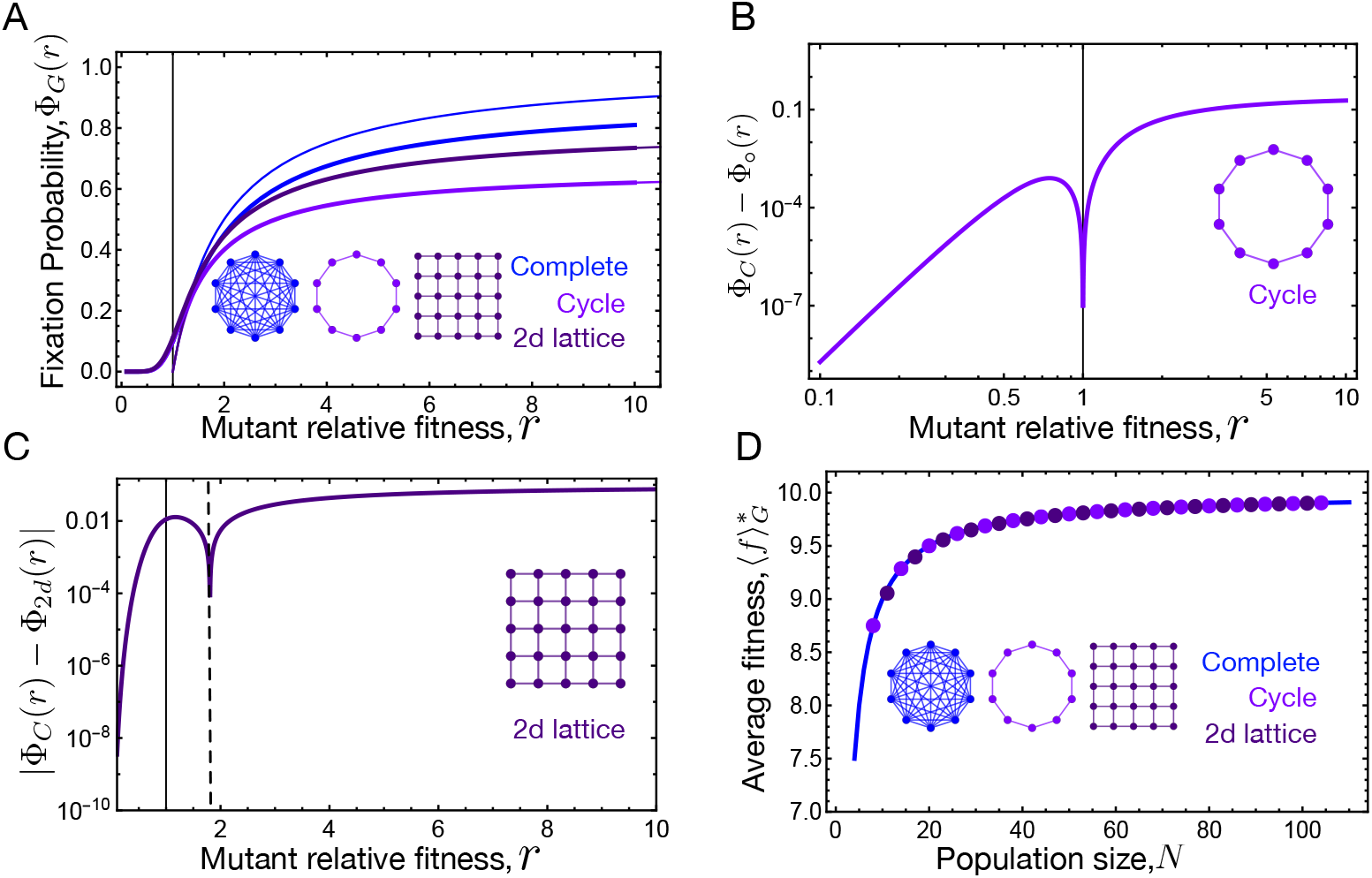
Regular graph: dB fixation probability and long-term evolution. A) Fixation probability for regular graphs – the cycle graph, the 2d lattice with periodic boundaries and the complete graph – under dB updating are shown. Thick lines correspond to population size *N* = 10. Thin lines represents lim_*N*→∞_ Φ_*G*_(*f′, f*). At fixation time scales, regular graphs behave differently. B) The cycle graph is a suppressor of fixation under dB updating. C) For small *N* (*N* =10 here), the 2d lattice behaves like a suppressor of selection. Φ_*C*_ − Φ_2*d*_ is negative for mutant fitnesses to the left of the dashed line, and positive otherwise. D) The cycle graph being a suppressor of fixation, attains the same steady-state average fitness in the mutation selection balance as the complete graph. For large *N*, 2*d* lattice attains the same fitness as the other graphs. At longer time scales, regular graphs have identical steady-state statistics contrary to their differences at shorter fixation time scales. Parameters: mutant fitness distribution, 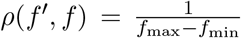 with *f*_min_ = 0.1 and *f*_max_ = 10.

The series D3 contains only odd terms because, by definition,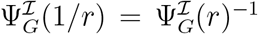. The second term of the series gives us the conservative estimate of the range of fitness values for which the origin-fixation dynamics is reversible, 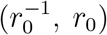 and *r*_0_ is found by setting a tolerance *ε*,

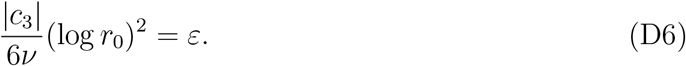

For the complete and the cycle graph, Ψ_*G*_ is a strict power-law with *c*_3_ = 0, which is why *r*_0_ → ∞ for these structures. However, for the 2d lattice, 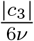 saturates to a finite value at larger *N*, see Fig. D.1, as both *c*_3_ and ν scales as *N* at large population sizes.

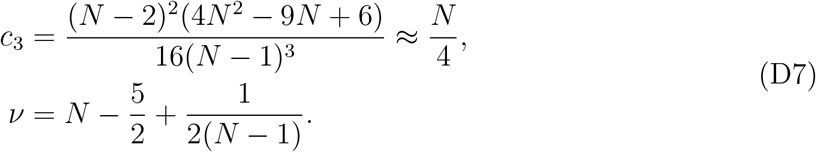

This makes *r*_0_ finite with a value approximately equal to 3 for ϵ = 0.05 and the range of fitness values where the Moran dB origin-fixation dynamics is reversible is *r* ∈ (0.33, 3). Above, we have used the relation,

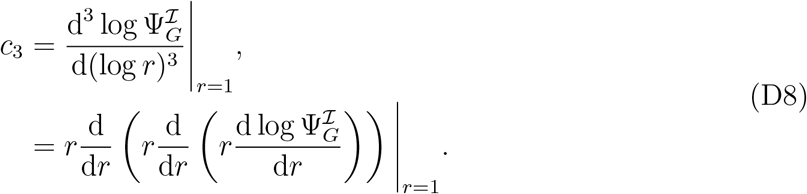

Similarly, 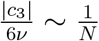 for the temperature initialised star graph under dB (dB^*p*^) dynamics. As a result *r*_0_ → ∞ in the limit *N* → ∞. The same result holds for the star graph under dB^*o*^ dynamics.

**FIG. D.1.**
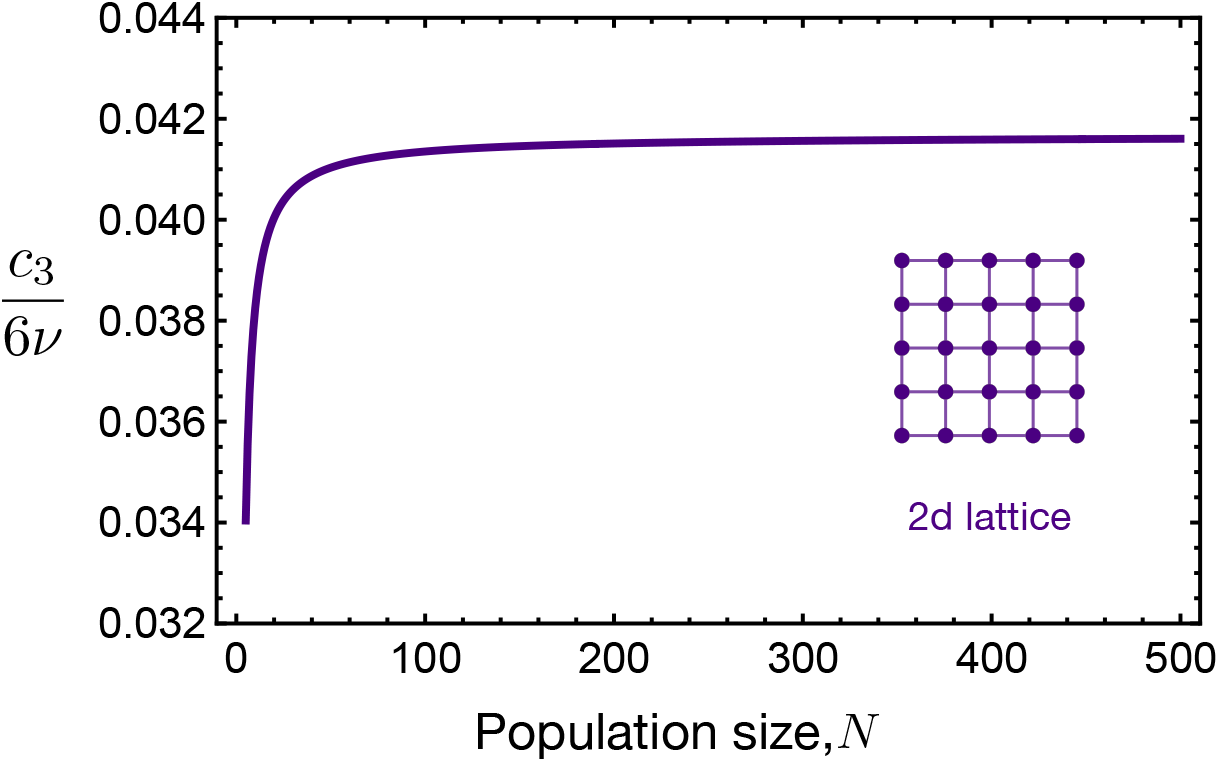
Reversible Moran dB origin-fixation dynamics on the 2d lattice. Coefficient 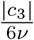 of the second term of Eq. D3 saturates to finite value for large *N*, which results in the range of fitness values where the origin-fixation dynamics is reversible to be *r* ∈ (0.33, 3).

## Appendix E

### Standard deviation in the steady-state fitness distribution

In this section, we derive the expressions for the standard deviation in fitness in the steady-state. For the complete graph we know that,

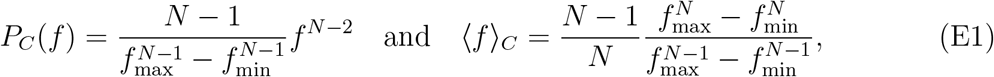

and

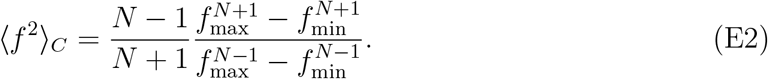

The variance turns out to be

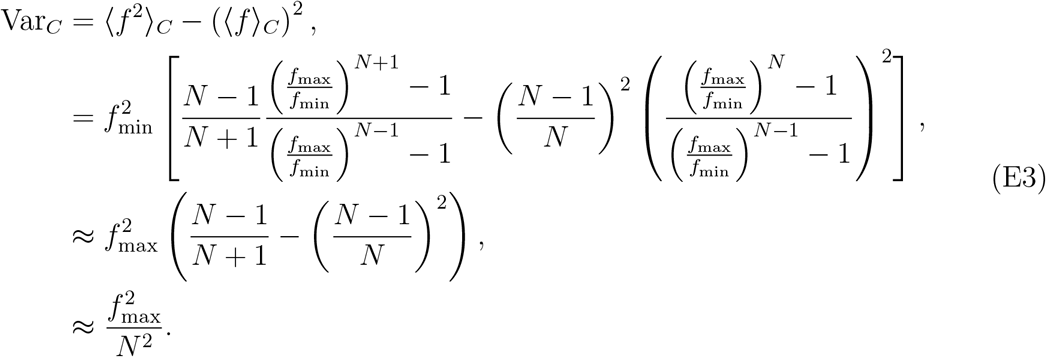

The above approximation holds for large population sizes. Therefore, the standard deviation in fitness for the complete scales as ∼1*/N*. For the star graph under dB^*o*^ dynamics, in the limit of large *N*

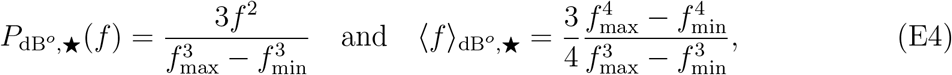

and

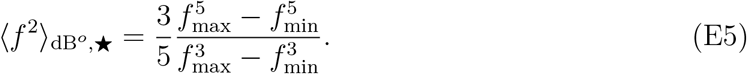

For the variance, we have

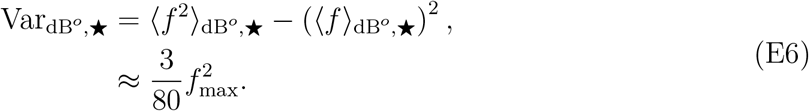

The standard deviation for the dB^*o*^ star graph is independent of *N* and asymptotes to a finite value. Similarly, for the dB^*p*^ star graph, for *N* 1 we have

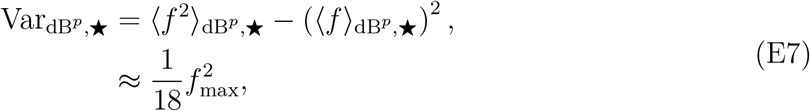

which is again independent of *N*. Note that for large *N*, 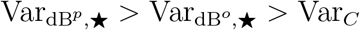. The order of fluctuations can be understood from the large *N* limit effective population size of the three structures −*N* for the complete graph, 2 for the dB^*o*^ star graph and 1 for the dB^*p*^ star graph.

Under Bd updating, we have seen that Ψ_*_ is same for both the Bd^*o*^ (Bd temperature initialisation) and Bd^*p*^ (Bd uniform initialisation) star graph. Consequently, the Bd^*o*^ and Bd^*p*^ have the same steady-state statistics and thus, variance in the steady-state,

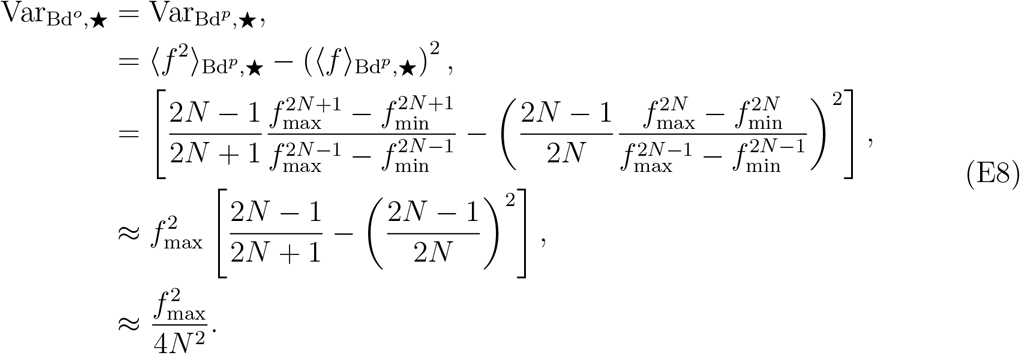

The above approximation holds for large *N*. Therefore, the standard deviation for the star graph under Moran origin-fixation Bd updating scales as 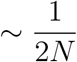.

## Appendix F

### Long-term evolution on discrete fitness space

To study Bd and dB long-term dynamics on random graphs we define a Markov chain. To proceed we first discretise the fitness space. The states are labelled with integer values from 0, 1, · · ·, *z*. For 1≤*i*≤*z* −1, the fitness of state *i* is equal to *f*_*i*_ = *f*_min_ + *i*·δ. The boundaries fitness are *f*_0_ = *f*_min_ and *f*_*z*_ = *f*_max_. The long-term dynamics on a graph *G* is a Markov chain obeying the equation,

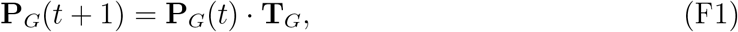

where **P**_*G*_(*t*) = (*P*_*G*,0_(*t*), *P*_*G*,1_(*t*), · · ·, *P*_*G,z*_(*t*)) with *P*_*G,i*_(*t*) being the probability for the population to be in fitness state *f*_*i*_ at time step *t*. The transition matrix **T**_*G*_ on the fitness space is given as,

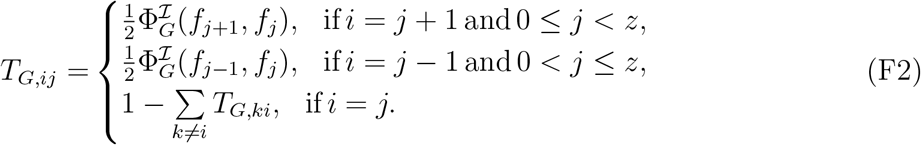

Matrix **T**_*G*_ is a positive and an irreducible matrix, therefore we can find the steady-state distribution 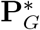 using the Perron-Frobenius theorem [92]. 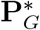 is the left eigenvector of the matrix **T**_*G*_ corresponding to eigenvalue 1,

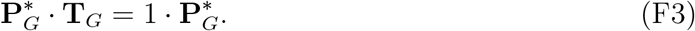

The steady-state average fitness is then,

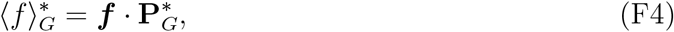

where ***f*** = (*f*_min_, *f*_1_, *f*_2_, *· · ·, f*_max_) is the fitness vector.

## Appendix G

### Criterion for a graph to have higher steady-state fitness than the complete graph

Just by looking at the 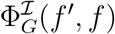, it is not easy to predict if the graph *G* will attain higher long-term average fitness than the complete graph in the mutation-selection balance of the Moran origin-fixation dynamics. The useful quantity for that purpose is the ratio of fixation probabilities, 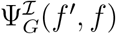 as defined in Eq. 12 (main text), which allows us to obtain a sufficient condition. Let us consider two graphs, *G*_1_ and *G*_2_, and denote the respective steady-state probability density functions for the Moran origin-fixation dynamics on these structures by 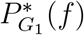 and 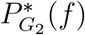.

**Proposition:** If the continuous density functions 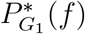 and 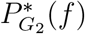 intersect exactly once, then

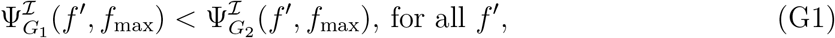

is a sufficient condition for 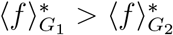.

**Proof:** From the condition (G1) and Eq. 12 of the main text, it follows that

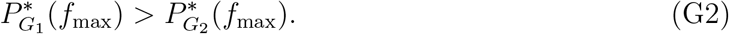

Using this fact, and denoting 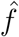 as the unique point at which the the functions intersect,

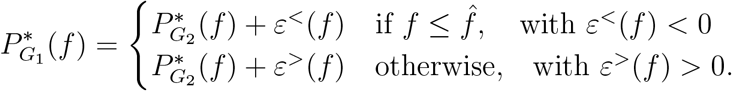

Then, from the normalization of the probability density functions, it follows that

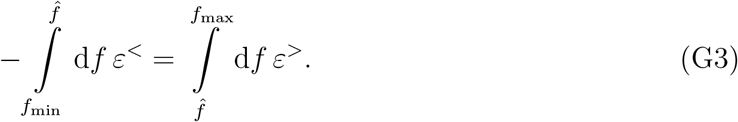

Now, note that

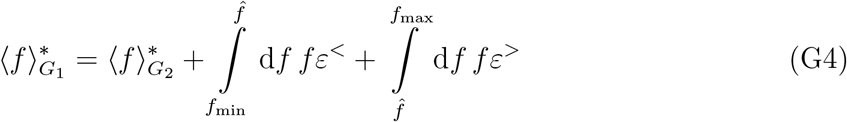

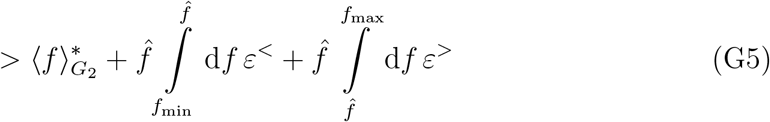

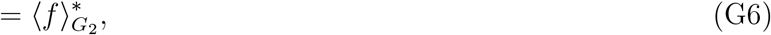

where the inequality follows straightforwardly from the signs of the integrands and the limits of the integrals, and the last equality follows from (G3). This completes the proof. □

The criterion 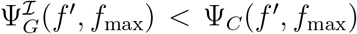 for all *f′*, is naturally satisfied by amplifiers of selection: It follows directly from the definition of amplifiers of selection, which are better at preventing the fixation of deleterious mutations and fixing advantageous mutants. Therefore, for an amplifier of selection,

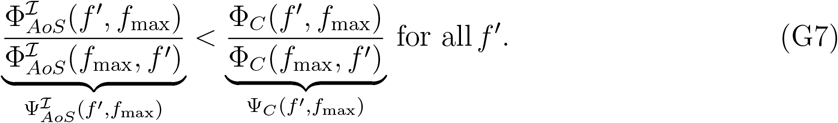

Subscript *AoS* denotes an amplifier of selection. Similarly, for a suppressor of selection,

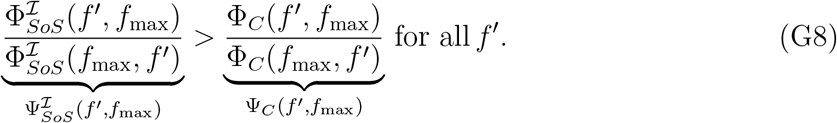

Subscript *SoS* denotes a suppressor of selection.

For suppressors of fixation, it is not obvious if they satisfy the criterion to attain higher average steady-state fitness in the mutation-selection balance. Suppressors of fixation have a lower probability of fixing beneficial mutations. As a result, 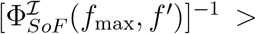(here *SoF* denotes a suppressor of fixation). However, they are also good in preventing the fixation of deleterious mutants,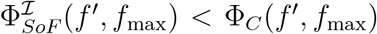. This means that a suppressor of fixation can satisfy the criterion G1 by compensating for its lower probability of fixation of beneficial mutants by rejecting deleterious mutants more efficiently. An example is presented in ref. [40]. Note that the criterion G1 holds for any update rule.

Let us apply the derived sufficient condition to the complete graph, dB^*o*^ and dB^*p*^ star graph. From Fig. G.1, we see that the steady-state probability density functions for any two graphs intersect at only one point, and thus we can use our criteria here. From Eqs. 14, 19, and 23, we have

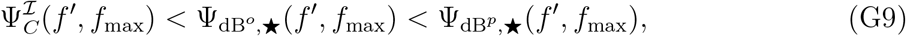

therefore, 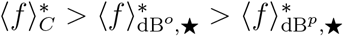.

**FIG. G.1.**
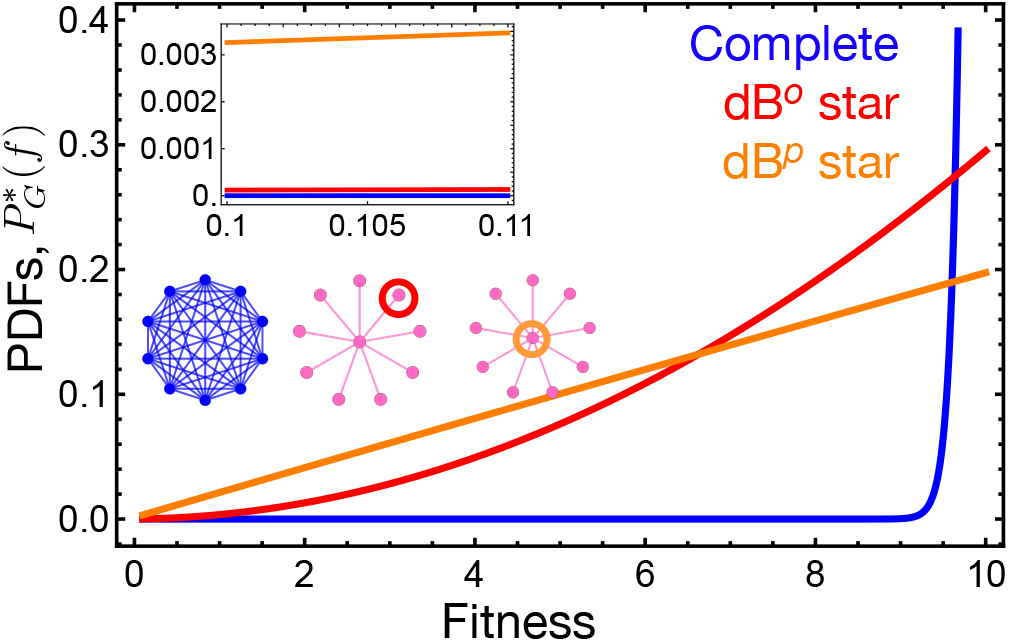
A pair of probability density functions intersect at one fitness point. The steady-state probability density functions (PDFs) for the complete graph, the dB^*o*^ and the dB^*p*^ star graph are shown. Firstly, the PDFs are monotonically increasing w.r.t fitness. Secondly, any pair of PDFs intersects at one point. Thus the sufficient condition G1 gives the ordering of average fitnesses. Parameters: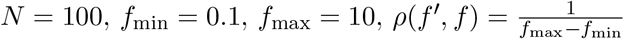.

## Appendix H

### Effects of deleterious mutants on the initial phase of the long-term dynamics

So far we have studied the effect of preventing/fixing deleterious mutants on the steadystate fitness statistics of graph. In this section, we explore the role of deleterious mutant regime on the initial phase of the mutation-selection dynamics, particularly the average selection coefficient of the first substitution given that the initial fitness is *f*,

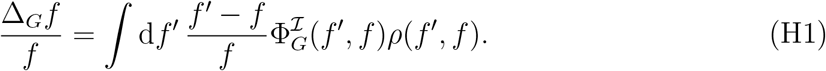

This quantity is also equal to the instantaneous rate of evolution [26], [25]. From Fig. H.1 A, we see that the mean selection coefficient decreases as the population is started with initial population fitness closer to the average steady-state fitness. The mean selection coefficient is anticipated to be positive if the initial fitness is below the average steady-state fitness and negative if started with higher fitness values. However, in Ref. [61], for the complete graph it has been shown that the mean selection coefficient of the first substitution is negative even if the population is initialised with fitness below the average steady-state fitness. We observe this effect to be more pronounced for the case of dB^*p*^ star graph. We investigate further by working out the large *N* case.

In the limit of *N*→ ∞, for the dB^*p*^ star graph under Moran dB^*o*^ updating and uniform mutational fitness distribution, we have

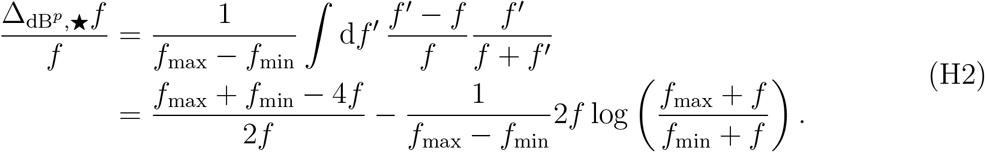

The value of *f* for which the mean selection coefficient (right hand side of the above equation) goes to zero is obtained numerically. The difference of the obtained fitness value and the average steady-state fitness for the dB^*p*^ graph, δ*f* is shown in Fig. H.1 B. For the parameters we use throughout the manuscript, the δ*f* for the dB^*p*^ star graph is finite, even in the limit of *N* → ∞. This means that if the long-term mutation-selection dynamics is initiated with fitness value in between 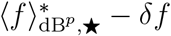 and 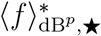, the first mutation to fix on average is deleterious. Therefore, the corresponding average fitness trajectories does not have monotonically increasing fitnesses. For the complete graph, in the limit of *N* → ∞,

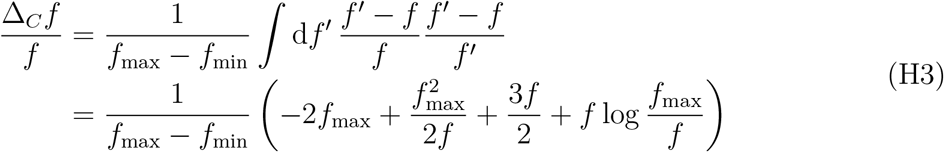

The r.h.s in the above equation goes to 0 for *f* = *f*_max_. Therefore, while δ*f* is non-zero for finite population sizes, in the limit of *N* → ∞, δ*f* = 0 for the complete graph.

**FIG. H.1.**
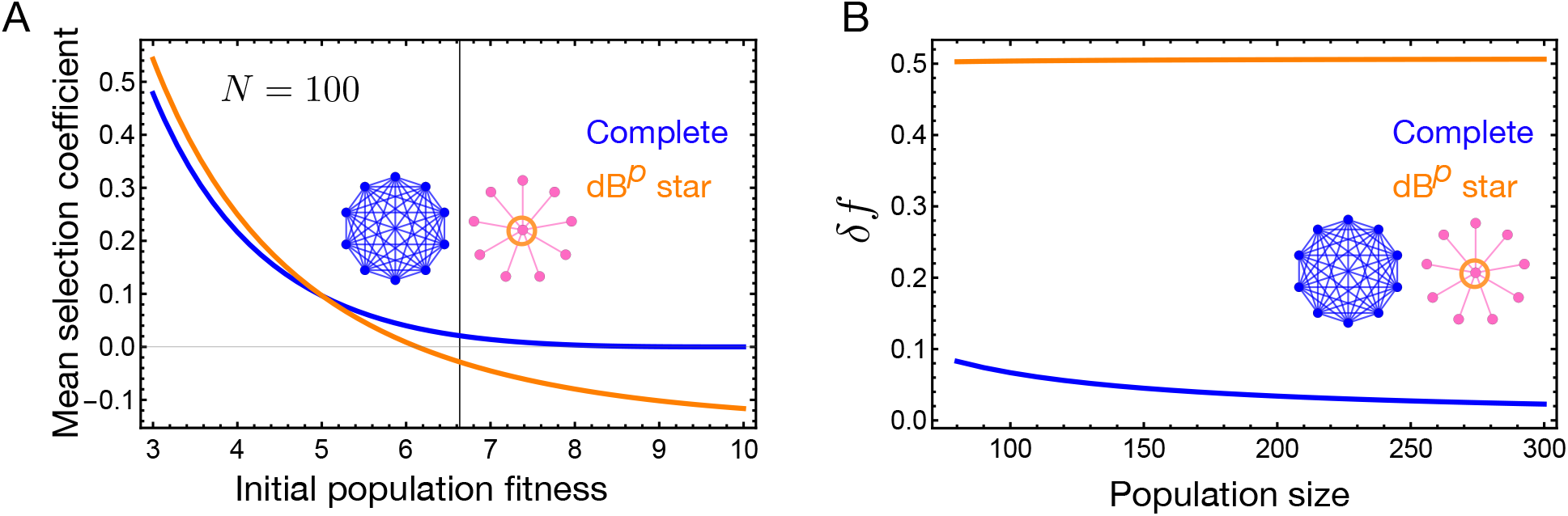
Mean selection coefficient of the first substitution. The black grid line in A) corresponds to the average steady-state fitness of the dB^*p*^ star graph. The mean selection coefficient is negative even if the population’s initial fitness is below the steady-state average fitness, thus leading to non-monotonic average fitness trajectories. B) Denoting δ*f* as the region of fitness values below the steady-state average fitness for which the mean selection coefficient is negative, in the limit of large *N* δ*f* asymptotes to 0.5 for the dB^*p*^ star graph, whereas the gap decays to 0 for the complete graph. Parameters: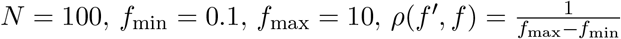.

## Appendix I

### Amplifier of fixation and suppressor of fixation in a metapopulation model

The amplifiers and suppressors of fixation are not solely restricted to one node one individual models, but can also be found in the network structured metapopulations [86]. The model in ref. [86] assumes time scale separation. The model has been analysed in the low migration rate regime where each node/deme is mainly in a monomorphic state. That is, when an individual migrates to a neighboring patch with a different type, either the migrating individual fixes or goes extinct before the next migration event occurs. The wild-type and the mutant type are assumed to have non-zero death rates, as a consequence of which the population sizes of fully mutant and wild-type demes are different. Here we assume that for both the individual types, the death rate is incorporated with the birth rate to give an effective growth rate. In this way, the population size for the mutant and wild-type deme is the same. The number of leaf demes is denoted by *d*. The probability that an individual from a given leaf deme migrates to the center deme is *m*_*I*_, whereas the probability that an individual from the central deme migrates to leaf deme is *m*_*O*_. The parameter *α* = *m*_*I*_*/m*_*O*_ quantifies the migration asymmetry. The probability that the mutant type with relative fitness *r* takes over the entire star network-structured metapopulation given that the population is initialised with central deme being the mutant deme and leaf demes being the wild-type demes is given by,

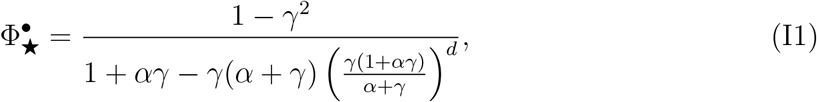

where

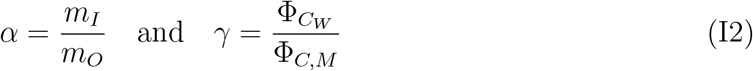

with

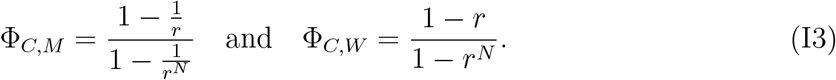

Assuming that the central deme is initialised as the mutant deme, we find that for smaller values of *α*, the star metapopulation acts as an amplifier of fixation. Conversely, for larger values of *α*, it functions as a suppressor of fixation, see Fig. I.1.

**FIG. I.1.**
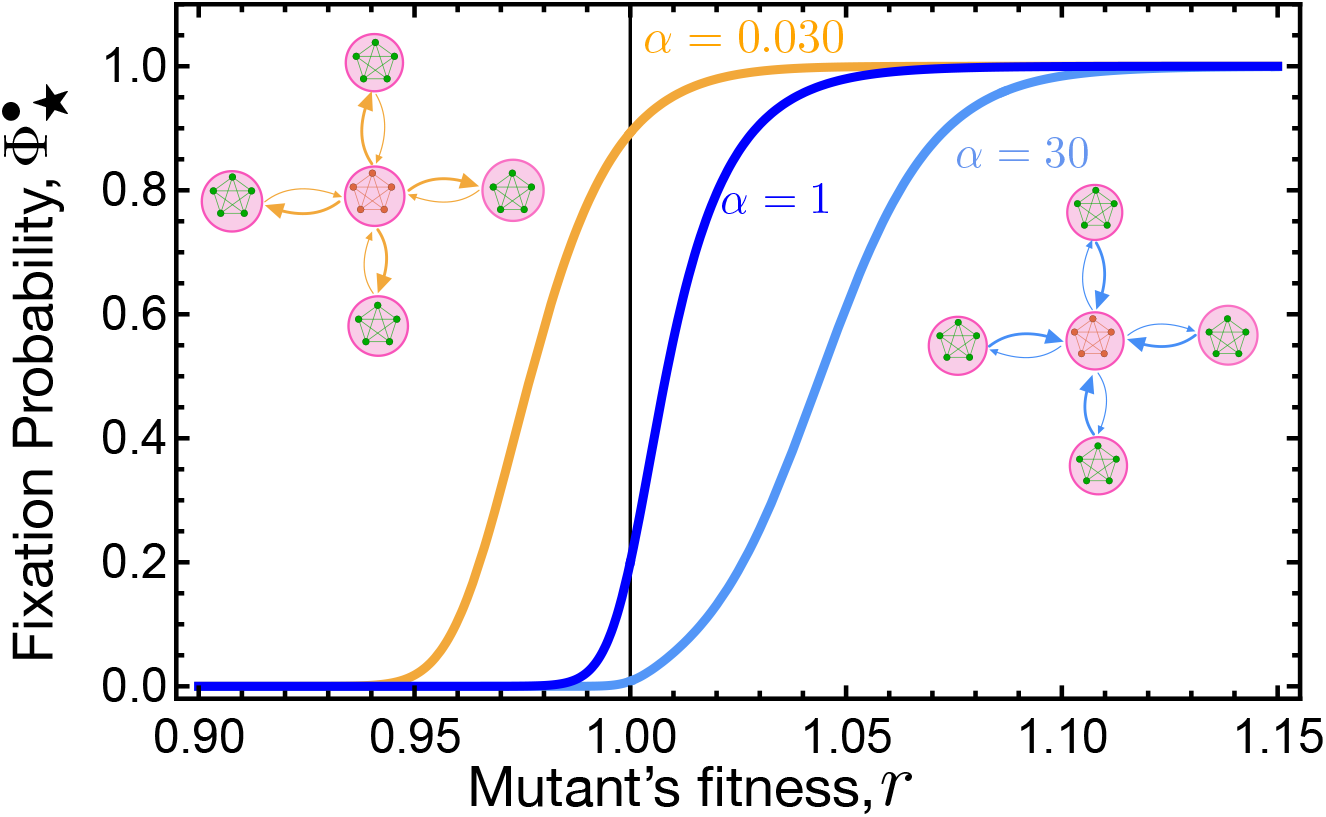
Amplifier and suppressor of fixation in meta-star. The probability that a mutant fixates in a star-structured metapopulation is shown given that the population is initialised with the central deme being the mutant deme. The probability that an individual from the central migrates to a leaf deme is *m*_*O*_, and the probability of migration for an individual migrating from a leaf node to the central node is *m*_*I*_. Parameter *α* = *m*_*I*_*/m*_*O*_ quantifies the migration asymmetry. According to the generalized circulation theorem in ref. [86], for *α* = 1, the meta-star’s fixation probability is the same as in the well-mixed population when started with *N* mutant individuals. Changing *α* from 1 gives two different fixation probability profiles: the amplifier of fixation (for lower *α*) and the suppressor of fixation (for higher *α*). We have encountered these profiles is the main text for one node one individual star under Bd^*o*^ and dB^*p*^. Parameters: size of each deme, *N* = 80 and the total number of demes *D* = *d* + 1 = 5.

